# Topology of gene regulatory compartments relative to the nuclear matrix

**DOI:** 10.1101/2022.05.10.491284

**Authors:** Gunhee Park, Kwangmin Ryu, Junho Lee, Geunhee Kim, Tae Lim Park, Hwanyong Shim, YigJi Lee, Changyeop Kim, Won-Ki Cho

## Abstract

Inside the nucleus, there are special compartments with characterized functions, some of which are involved in gene expression^1^. These compartments include transcriptional condensates and nuclear speckles, which contain factors required for transcription and splicing, respectively. While the characteristics of these intranuclear compartments were extensively investigated, spatial relationship between them is yet unclear. Meanwhile, RNA-protein structural network named the nuclear matrix has been observed inside the nucleus and suggested as a framework for the spatial segregation of the intranuclear space^2–6^. However, concept of the nuclear matrix was not widely accepted due to its lack of in vivo evidence^7^. Here, we report visualization of the nuclear matrix and intranuclear compartments using super-resolution fluorescence microscopy. We found the nuclear matrix is dynamic in live cells and easily disrupted upon transcription inhibition. Remarkably, we observed an orderly layered distribution of transcriptional condensates and nuclear speckles relative to the nuclear matrix. We also observed a separation of chromosome territories from transcriptional condensates and nuclear speckles by the nuclear matrix. Based on our findings, we propose a topological model for the regulation of transcription across the nuclear matrix.

## Introduction

Timely expression of appropriate levels of specific genes in response to external stimuli or differentiation signals should require a precise organization of the gene expression machinery in the nucleus. It is clear that nuclear chromatin is compartmentalized in chromosome territories and that various specialized subnuclear organelles are involved in the regulation of gene expression^1^. It remains unclear how these subnuclear compartments are organized and how the spatial framework that coordinates their localization and function works.

In the mid-1970’s, a mesh-like structural network partitioning the intranuclear space was visually discovered via electron microscopy and dubbed the nuclear matrix^2, 7^. In these experiments, cells were subjected to a series of sample preparation steps to isolate an insoluble nuclear fraction that included RNA and proteins. Since the earliest observation, specific components of the nuclear matrix have been identified, including the abundant nuclear protein scaffold attachment factor A (SAF-A), which is also known as heterogeneous nuclear ribonucleoprotein U (hnRNPU)^8, 9^.

SAF-A was first known to be involved in pre-mRNA splicing in the nucleus^10, 11^. While SAF-A is an RNA-associated ribonucleoprotein, it also associates with specific DNA regions called scaffold/matrix attachment regions^12^. In addition to potentially contributing to the formation of the nuclear matrix, SAF-A oligomerizes *in vitro*^13^. Because most of these observations were accomplished via electron microscopy or biochemical assays that required harsh sample preparation protocols, we asked whether the nuclear matrix exists and can be visualized in living cells. Contingent on successful visualization of the nuclear matrix in living cells and because SAF-A associates with both DNA and RNA while promoting RNA processing, we hoped to visualize any spatial correlations between chromosome territories and gene expression-related subnuclear compartments, transcriptional condensates, or nuclear speckles.

## Results

### Visualization of RNA-dependent nuclear matrix *in vivo*

To visualize the nuclear matrix in live cell nuclei, we used a CRISPR/Cas9-mediated knock-in technique^14^ to label the C-terminus of endogenous SAF-A in the human colon cancer cell line HCT116 with Dendra2, a photoconvertible fluorescent protein for stochastic localization-based super-resolution imaging (Fig. 1, Extended Data Fig. 1). Since SAF-A is one of the most abundant nuclear proteins^9^, conventional fluorescence imaging did not allow us to resolve any detailed subnuclear structures (Fig. 1a). With super-resolution microscopy, however, we were able to observe a SAF-A structural network *in vivo* in a two-dimensional focal plane that was similar in appearance to the images of the nuclear matrix previously captured by electron microscopy^3–6^ (Fig. 1b-e). We also visualized similar nuclear matrix structures by labeling endogenous SAF-A with the far-red organic dye JF646 via a HaloTag in HCT116 cells (Extended Data Fig. 1, 2) and in human retinal pigment epithelial-1 (hTERT-RPE1) cells (Extended Data Fig. 2). SAF-A disassociates from chromosomes as they enter mitosis^3^. We confirmed that SAF-A was excluded from chromosomes in our endogenously labeled cell lines as they entered mitosis, ruling out the possibility that fluorescence protein-tagging of SAF-A prevents it from detaching from mitotic chromosomes (Extended Data Fig. 3). These results suggest fluorescently tagged endogenous SAF-A retains its localization and function in live cell nuclei.

**Fig. 1.**
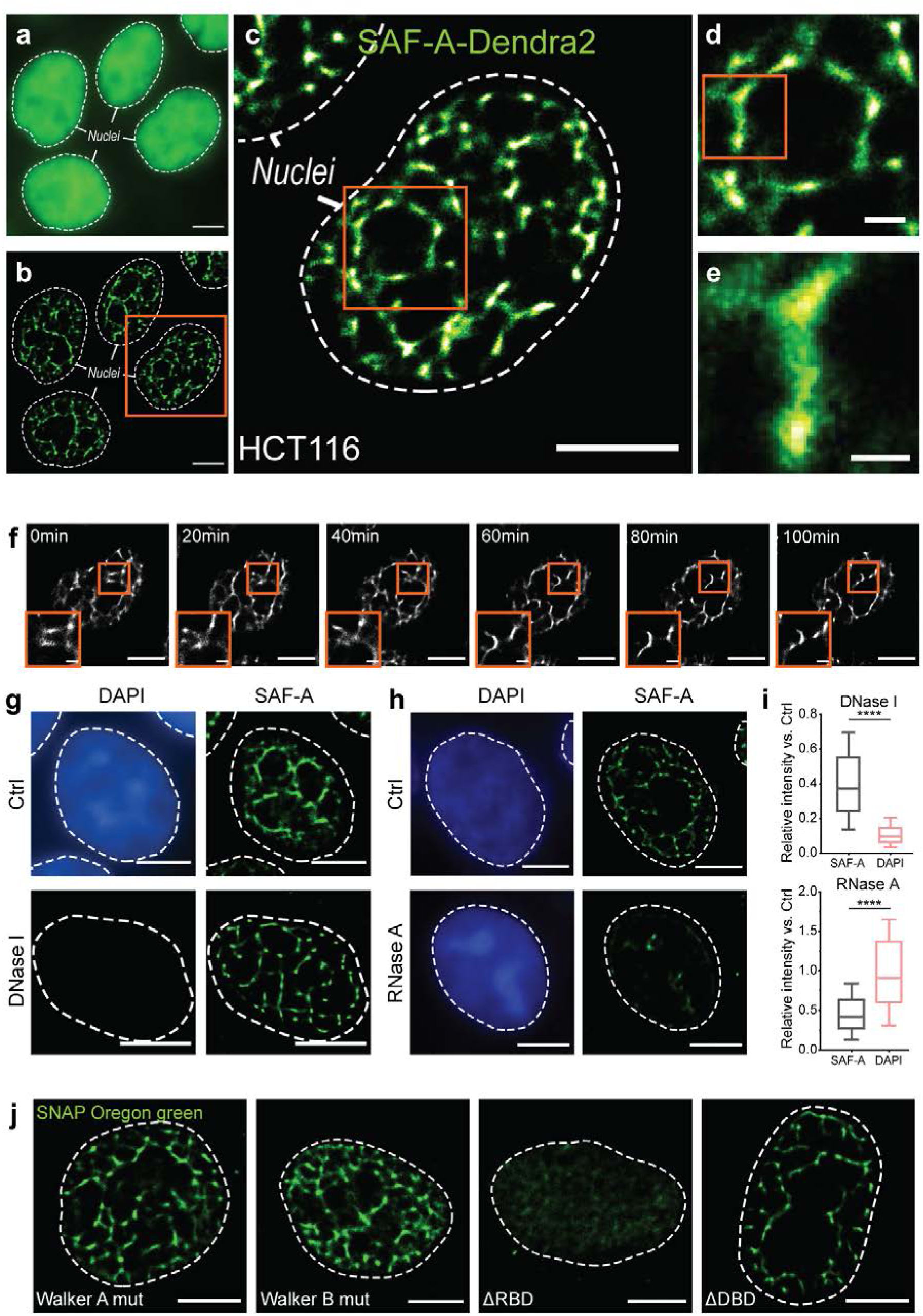
Labeled endogenous SAF-A forms dynamic nuclear matrix structures in living cell nuclei. **a, b**, Conventional (**a**) and super-resolved fluorescence images (**b**) of Dendra2-labeled SAF-A in HCT116 cell nuclei. **c-e**, Representative super-resolved images. Successive images are magnified versions of the boxed area in the previous image. SAF-A forms a structural network that contributes to the nuclear matrix. **f**, Time-lapse super-resolved images show dynamic changes in the nuclear matrix. **g, h**, Representative super-resolved images after treatment with DNase I and RNase A, respectively. **i**, Relative intensity of SAF-A-Dendra2 and DAPI after DNA or RNA degradation. **j**, Representative super-resolved images of the nuclear matrix in cells overexpressing Dendra2-labeled SAF-A mutants for the indicated domains after precise degradation of endogenous SAF-A. Scale bars represent 5 µm for entire nuclei images and 1 µm or 500 nm for zoom-in images.

When we performed a volumetric super-resolution imaging experiment, we confirmed that the structural network of the nuclear matrix extends throughout the entire nucleus (Extended Data movie S1). Moreover, we found via time-lapse super-resolution imaging experiments that the structure of the nuclear matrix changes dynamically through interphase (Fig. 1f, Extended Data Fig. 4, and movie S2).

We next asked whether the structures we observed with super-resolution fluorescence imaging consistently show the RNA-dependent insolubility that previous studies have assigned to the nuclear matrix^6^. To remove the soluble parts of the nucleus, we subjected our cell lines to an *in situ* fractionation protocol that included an extraction of nuclear components with the surfactant Triton X-100 and a subsequent DNA (DNase I) or RNA (RNase A) degradation step^6^. We found that although DNA degradation had only minor effects, RNA degradation dramatically disrupted the structure of the network, indicating that the nuclear matrix is RNA-dependent (Fig. 1g, i, Extended Data Fig. 5). We additionally observed an enrichment of nascent RNAs in the nuclear matrix (Extended Data Fig. 6). We noted that DNA degradation slightly reduced the intensity of the SAF-A signal across the nucleus despite the nuclear matrix structure remaining intact. This suggests the presence of some SAF-A molecules in the soluble fraction^13^ (Fig. 1i).

From its N- to C-termini, SAF-A structurally consists of DNA binding (SAP), ATP binding and hydrolysis (AAA+), and RNA binding (RGG) domains^13, 15^. To determine which of these domains are required for the formation of the nuclear matrix, we repeated our imaging experiments with cells overexpressing SAF-A mutants lacking each domain, followed by acute depletion of endogenous SAF-A via the auxin-induced degron system^16^ (Fig. 1j, Extended Data Fig. 7). We found mutants lacking the RNA binding domain failed to form the nuclear matrix structure, suggesting SAF-A binding to RNA is required for nuclear matrix formation. In contrast, neither the ATP binding and hydrolysis domain nor the DNA binding domain were essential for nuclear matrix formation.

Consistent with a previous study, we next found via dual-color imaging of SAF-A with immunolabeled representative epigenetic modifications (i.e., H3K27ac and H3K27me3) that the nuclear matrix is depleted from transcriptionally inactive heterochromatin regions and enriched in transcriptionally active euchromatin regions (Extended Data Fig. 8)^12^. Together, our results indicate that the formation and maintenance of the nuclear matrix is tightly linked to transcription.

### Coordinated distribution of transcriptional condensates around the nuclear matrix

We next sought to investigate the intranuclear positions of transcriptional condensates relative to the nuclear matrix using dual-color super-resolution imaging. Transcriptional condensates comprising RNA Polymerase II (Pol II), Mediator, and their coactivators are associated with active genes^17–19^. Other studies demonstrated that SAF-A-bound RNAs are enriched for intron sequences^11, 20^. These parallel observations, together with our own results described above, imply a spatial relationship between the nuclear matrix and transcriptional condensates. To visualize both structures simultaneously in live nuclei, we generated an HCT116 cell line in which endogenous SAF-A and Pol II are tagged with Dendra2 and HaloTag, respectively^21^ (Extended Data Fig. 1).

Super-resolution imaging of SAF-A and Pol II revealed a close apposition of the nuclear matrix and transcriptional condensates (Fig. 2a). It is interesting to note that the transcriptional condensates we observed were adjacent to the nuclear matrix, but not co-localized with it. Furthermore, most of the transcriptional condensates were positioned along the inner boundary of spaces defined by the nuclear matrix.

**Fig. 2.**
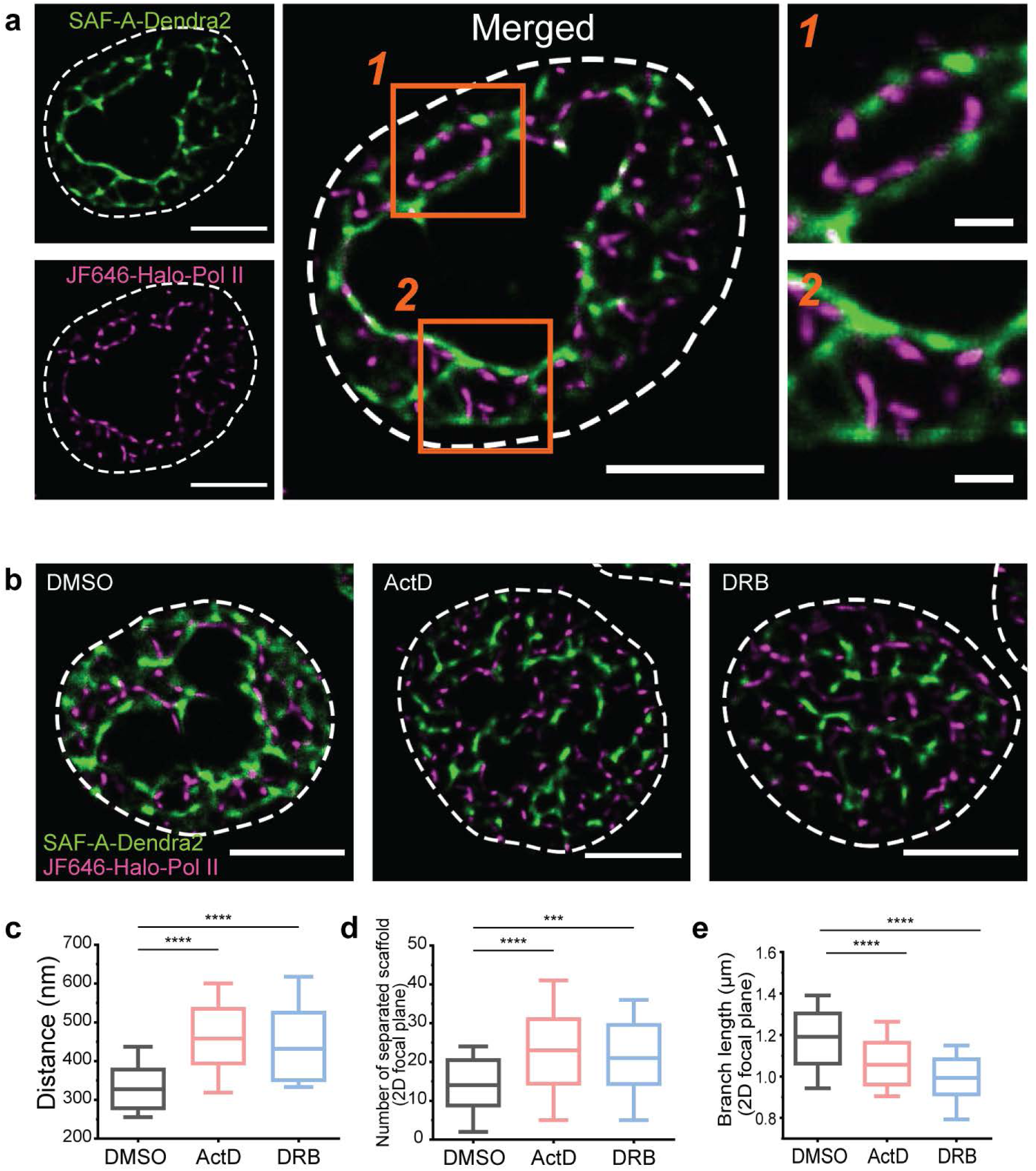
Transcriptional condensates appear along the inner boundary of spaces defined by the nuclear matrix. **a**, Dual-color super-resolution imaging of SAF-A-Dendra2 (green) and JF646-Halo-Pol II (magenta). Magnified images show the relative positions of transcriptional condensates and the nuclear matrix. **b**, Dual-color images of transcriptional condensates and nuclear matrix in the presence of the transcription inhibitors, Actinomycin D or DRB. **c-e** The distance between transcriptional condensates and the nearest nuclear matrix signal (**c**), the number of distinct nuclear matrix signal foci (**d**), and the length of continuous nuclear matrix network (**e**) as quantified via a skeletonization analysis before and after transcription inhibition. Scale bars represent 5 µm for entire nuclei images and 1 µm for zoom-in images.

Because the nuclear matrix is RNA-dependent and because of the coordinated positioning of the nuclear matrix and transcriptional condensates, we hypothesized that the structure of the nuclear matrix depends on transcription. When we visualized the nuclear matrix in the presence of the transcription inhibitors actinomycin D (ActD) or 5,6-Dichlorobenzimidazole 1-β-D-ribofuranoside (DRB), we observed a dramatic disruption of the nuclear matrix structure (Fig. 2b, Extended Data Fig. 9). Remarkably, transcription inhibition also led to an uncoupling of the position of the nuclear matrix and transcriptional condensates. To quantify these changes, we used a skeletonization analysis to define the structure of the nuclear matrix^22, 23^ and applied a density-based spatial clustering of applications with noise (DBSCAN) algorithm to the transcriptional condensates^24, 25^ (Fig. 2c-e, Extended Data Fig. 10). The average minimum distance between the transcriptional condensates and the skeletonized nuclear matrix increased significantly upon transcription inhibition (Fig. 2c) because of nuclear matrix fragmentation (Fig. 2d, e, Extended Data Fig. 10). Furthermore, by washing out the DRB and allowing transcription to resume, we were able to restore the spatial relationship between the nuclear matrix and the transcriptional condensates (Extended Data Fig. 11). The rapid redistribution of SAF-A upon transcription inhibition and restoration via DRB removal suggests the nuclear matrix is a highly dynamic structure and that newly transcribed RNAs contribute to its formation. In addition, we note that DRB induced a redistribution of SAF-A, radially concentrating it toward the center of the nucleus (Fig. 2b, right, Extended Data Fig. 12).

### Layered structure of gene regulatory compartments relative to the nuclear matrix

Based on these results, we next asked whether nuclear speckles, which have been implicated in splicing, are also spatially related to transcriptional condensates and nuclear matrix. A spatial correlation seems likely because the splicing of nascent RNAs is coupled to transcription and because previous studies have demonstrated an association between intronic RNAs and SAF-A^11, 26, 27^. Thus, we visualized nuclear speckles by immunolabeling the nuclear speckle component and splicing factor SC35 in cells with endogenously labeled SAF-A and Pol II (Fig. 3).

**Fig. 3.**
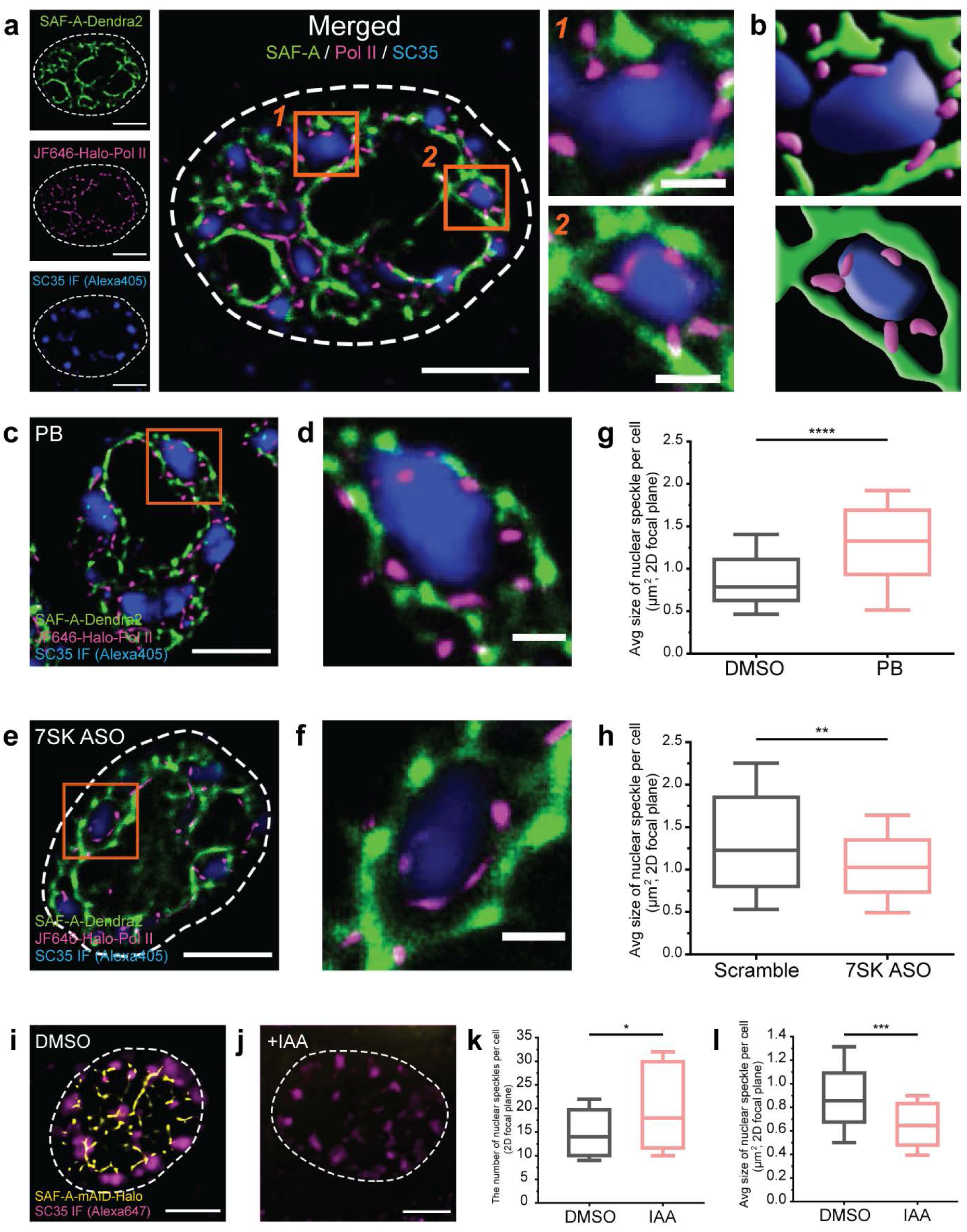
Nuclear speckles and transcriptional condensates form a layered structure within spaces defined by the nuclear matrix. **a**, Visualizing nuclear speckles via immunolabeling SC35 with Alexa405 (blue), nuclear matrix (SAF-A-Dendra2, green), and transcriptional condensates (JF646-HaloTag Pol II). Magnified images clearly show a radially layered structure in which nuclear speckles, transcriptional condensates, and nuclear matrix are arranged in order from the inside toward the outside. **b**, Cartoonization of the magnified images in (A) based on fluorescent signal contours and intensities. **c-f**, Visualization of nuclear speckles, transcriptional condensates, and nuclear matrix after PB-induced splicing inhibition (**c, d**) or ASO-induced depletion of 7SK lncRNA (**e, f**). **g, h** Average nuclear speckle size before and after PB treatment or 7SK depletion, respectively. **i-l**, Representative images of nuclear speckles (**i, j**), plots of the number of nuclear speckles per cell (**k**), and plots of average nuclear speckle size before and after acute SAF-A depletion (**l**). Scale bars represent 5 µm for entire nuclei images and 1 µm for zoom-in images.

In these experiments, we observed nuclear speckles as condensed regions of SC35 signal that appeared alongside the nuclear matrix and transcriptional condensate signals (Fig. 3a). To our surprise, nuclear speckles, transcriptional condensates, and the nuclear matrix exhibited a layered structure in which the nuclear speckles are positioned in the center of spaces defined by the nuclear matrix (Fig. 3a, b). Remarkably, transcriptional condensates appeared around each nuclear speckle. This result seems consistent with the hypothesis that nuclear speckles are sites of splicing factor storage rather than active RNA splicing zones^28, 29^. This would mean that the machinery necessary for RNA splicing is immediately accessible to sites of active gene expression. Consistent with our observations and with previous studies showing that poly-adenylated RNAs co-localize with nuclear speckles^30, 31^, we propose a model in which mature RNAs are transported to nuclear speckles after transcription and spliced introns are transported in the opposite direction to participate in the formation of the nuclear matrix (Fig. 4c)^11, 26, 27^.

**Fig. 4.**
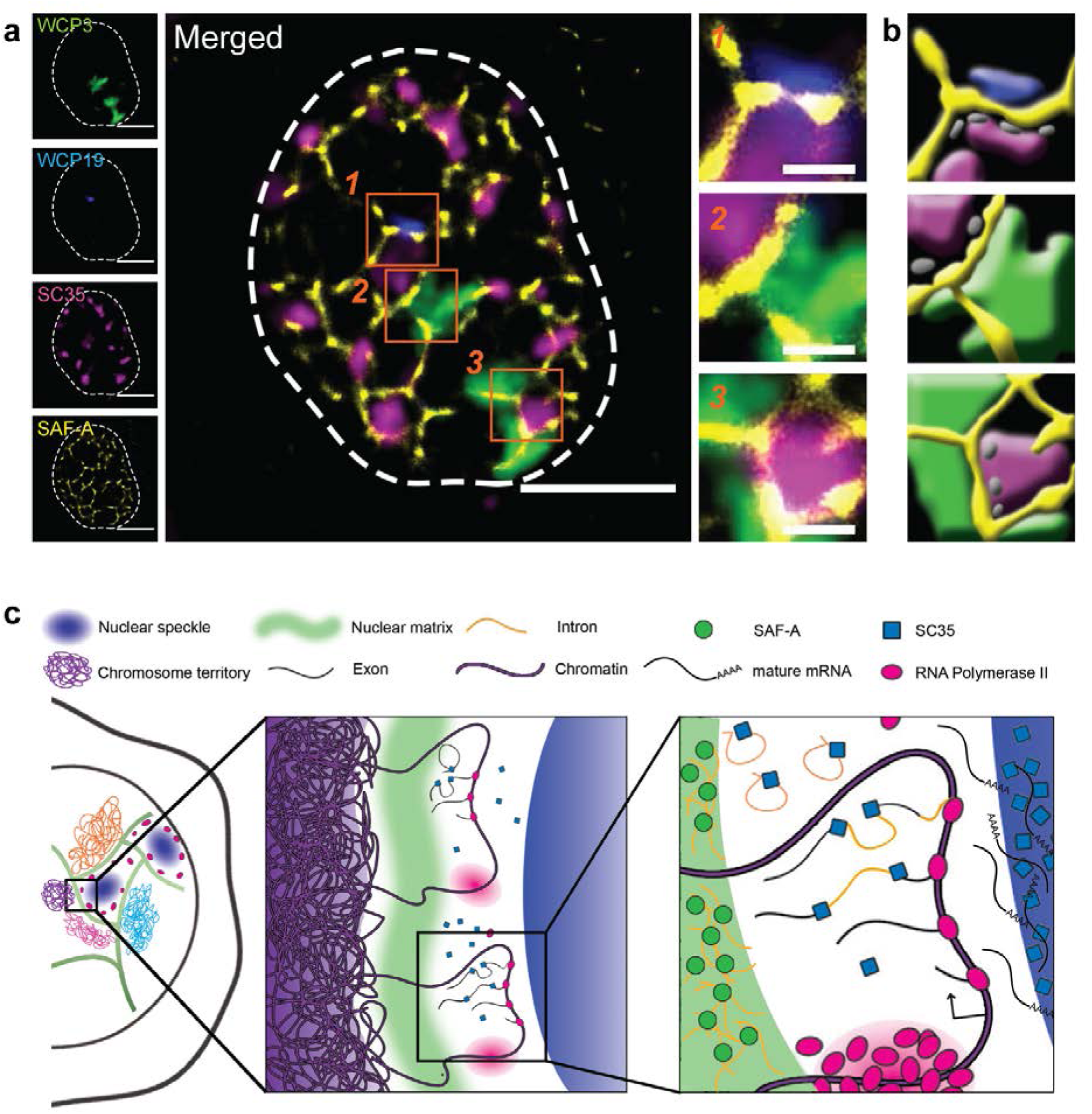
Chromosome territories remain separate from transcriptional condensates and nuclear speckles across the nuclear matrix. **a**, Mapping chromosome territories (Chromosome 3, green, and Chromosome 19, blue) in relation to the nuclear matrix (SAF-A-HaloTag-JF646, yellow) and nuclear speckles (SC35 immunolabeling with Alexa405, magenta). Magnified images clearly distinguish the separation of nuclear speckles from chromosome territories by the nuclear matrix. **b**, Cartoonization of the magnified images in **a** based on fluorescent signal contours and intensities. Transcriptional condensates shown in gray are hypothetically added. Scale bars represent 5 µm for entire nuclei images and 1 µm for zoom-in images. **c**, A schematic representation of our topological gene expression regulation model functioning across the nuclear matrix.

To determine whether the nuclear matrix and nuclear speckles regulate one another, we perturbed nuclear speckles by depleting the 7SK lncRNA via transfection with antisense oligos (ASO) or by inhibiting splicing with Pladienolide B (PB) (Fig. 3c-f, Extended Data Fig. 13). Upon PB-induced splicing inhibition, we observed an increase in average nuclear speckle size (Fig. 3g). In contrast, 7SK depletion via ASO reduced average nuclear speckle size (Fig. 3h). Interestingly, the layered radial organization—nuclear speckle to transcriptional condensate to nuclear matrix in the outward direction—was maintained under both conditions (Fig. 3d, f). DRB-induced transcription inhibition, however, significantly disrupted this layered structure, decoupling nuclear speckles from transcriptional condensates with the disruption of the nuclear matrix (Extended Data Fig. 14). These results indicate not only that the layered structure we observed is transcription-dependent, but also that the nuclear matrix is flexible toward nuclear speckles rather than physically restricting them as a solid barrier.

In addition, we acutely depleted SAF-A using the auxin-induced degron system and asked whether nuclear speckles were affected (Fig. 3i, j, Extended Data Fig. 1). We found loss of SAF-A increases the number of nuclear speckles but reduces average sizes (Fig. 3k-l). These results imply that the nuclear matrix does not simply form near nuclear speckles in a passive manner; it also influences nuclear speckle formation.

### Nuclear matrix segregates chromosome territories from gene regulatory compartments

We next performed whole chromosome painting (WCP) with fluorescence in situ hybridization (FISH) probes^32, 33^ to visualize chromosome territories relative to the nuclear matrix (Fig. 4a, b, Extended Data Fig. 15). As expected, the resulting WCP images showed either one or two of each painted chromosome at a fixed focal plane. Interestingly, we found that the chromosome territories and the nuclear matrix did not overlap. We also found that the WCP probes showed weak labeling at the chromosome territory borders in what are presumably regions of gene-rich euchromatin.

Mapping the chromosome territories relative to the nuclear matrix and nuclear speckles indicated, to our surprise, that the chromosome territories were separated from the nuclear speckles by the nuclear matrix (Fig. 4a, b, Extended Data Fig. 15). We noted that the borders of the chromosome territories contact nuclear speckles across the nuclear matrix. In addition, by labeling the nucleolar marker fibrillarin, we confirmed that nucleoli are also separate from the nuclear matrix (Extended Data Fig. 16).

## Discussion

Integrating these observations together, we suggest a model for the topologically specialized regulation of transcription relative to the nuclear matrix. In this model, a layered structure forms in the nucleus, with chromosome territories, nuclear matrix, transcriptional condensates, and nuclear speckles appearing in a stereotyped order (**Fig. 4c**). In this model, regions of euchromatin loop out from the chromosome territories across the nuclear matrix so that transcription can take place at active gene loci near the edges of nuclear speckles, where splicing processes can be facilitated. This model supports the concept that interchromosomal ‘active hub’ is formed around the nuclear speckles, suggested by recent split-and-pool sequencing studies^34^.

SAF-A and RNA each are known to contribute to the maintenance of euchromatin^12, 13^. Our observation, putting SAF-A into the context of the nuclear matrix provides an integrated view of how transcription-dependent RNA-protein network affects the chromatin state. The nuclear matrix, which is composed of SAF-A and RNAs, presumably mostly introns that have been spliced from active gene transcripts, may hold active genes around nuclear speckles, or segregate active genes from condensed chromatin, to maintain active gene regions in their decondensed states^15^.

## Methods

### Cell lines

HCT116 CMV-OsTIR1 cells from a previous study^21^ and all CRISPR/Cas9 knock-in cell lines derived from them were cultured at 37°C in 5% CO2 in DMEM (HyClone) supplemented with 10% FBS (Gibco) and 100U/ml penicillin and 100μg/ml streptomycin (HyClone). Cell lines were tested for Mycoplasma contamination using the e-MycoTM Mycoplasma PCR Detection kit (iNtRON).

### Homology-directed repair (HDR) DNA template design

To generate SAF-A C-terminal Dendra2 knock-in repair template, 500bp of left homology arm and right homology arm containing overhang were amplified from gDNA of HCT116 cells by using Q5 High Fidelity DNA Polymerase (NEB, M0491). The silent mutation was included in the repair template to avoid cleavage by Cas9. Double BbsI restriction sites flanking homology arm pair fragments were amplified by overlap extension PCR and inserted at the multi-cloning site of the pUC19 vector. The Dendra2 sequence (690bp) was inserted at BbsI-HF (NEB, R3539) digested pUC19-SAF-A C-term repair template backbone vector. In the case of the SAF-A C-term mini-AID-Halo knock-in repair template, the mAID-Halo construct (1101bp) was cloned into the repair template backbone vector.

To generate RPB1 N-term HaloTag knock-in repair template, 500bp of homology arm pair which is flanking double BsaI restriction site and containing GS-linker at upstream of 3’ homology arm, was synthesized by Gene Synthesis (Bionics, Seoul, South Korea) and inserted at the multi-cloning site of the pUC57 vector. The silent mutations was included in the repair template to avoid cleavage by Cas9. The HaloTag sequence (891bp) was inserted at BsaI-HF®v2 (NEB, R3733) digested pUC57-RPB1 N-term repair template backbone vector.

### CRISPR/Cas9 engineering for endogenous SAF-A and Pol II labeling

Single guide RNAs (sgRNAs) targeting 50bp upstream or downstream from the stop codon of SAF-A or start codon of RPB1 were designed using the CRISPR Design tool (https://sg.idtdna.com/site/order/designtool/index/CRISPR_CUSTOM).

DNA oligonucleotides for sgRNA were obtained from Bioneer (Daejeon, South Korea) and annealed in an annealing buffer. Annealed oligonucleotide fragments were inserted at Streptococcus pyogenes Cas9 vectors (pSpCas9 (BB)-2A-Puro (PX459) V2.0, Addgene #62988) which are digested by BbsI-HF enzyme. Each 1μg of sgRNA-Cas9 plasmid and repair template plasmids were transfected to 5x105-1x106 cells by FuGene HD (Promega, Cat#2311) according to the manufacturer’s protocol.

More than 72 hr after co-transfection of sgRNA-Cas9 plasmids and repair templates, the transfected cells were sorted by FACS (Bio-rad, S3e cell sorter) to establish knock-in cell lines. For knock-in cells harboring Dendra2, 488-nm illumination was used for sorting. For knock-in cells harboring HaloTag, 640-nm illumination was used after 15 min incubation with 100nM Janelia Fluor® HaloTag Ligands (JF646 HaloTag, Promega, GA1120). Fluorescence-positive cells were sorted and collected into cell growth media and grown in 48 well cell culture plates. Successful CRISPR/Cas9 gene editing of the knock-in cell lines were confirmed by genomic DNA PCR and fluorescence microscopy.

To generate monoclonal homozygous SAF-A C-term mAID-Halo knock-in cell line, fluorescence-positive single-cells were sorted into 96-well cell culture plates. The genotype of each monoclonal cell line was confirmed with genomic DNA PCR.

### PCR genotyping

Genomic DNAs of each cell lines were harvested by gDNA extraction kit (Enzynomics) according to the manufacturer’s instruction. Primer pairs targeting boundaries of the 5′ and 3′ knock-in sites were designed to confirm the insertion of the Dendra2 or mAID-Halo sequence at the target site (Supplementary Table S1). Target DNA for genotyping was amplified using 2XTOPsimple™ DyeMix-nTaq (Enzynomics) or Q5 High fidelity DNA polymerase (NEB Cat#M0491) with initial denaturation at 95°C for 5 min, followed by 30 cycles of denaturation at 95°C for 10 s, annealing at 60°C for 30 s, extension at 72°C for 1 to 2 min, and a final elongation step at 72°C for 5 min.

### In situ fractionation, RNase/DNase treatment

HCT116 cells harboring endogenous SAF-A-Dendra2 were grown on Poly-L-ornithine (PLO) coated glass bottom dish. For dual-color imaging of H2B and SAF-A, cells were transfected with expression vector harboring H2B-EBFP2 (Addgene #55243) using FuGene HD. For in situ fractionation, cells were permeabilized with 0.5% Triton X-100 in cytoskeleton (CSK) buffer for 3 min on ice, then treated with RNase A or DNase I as described below.

For RNase treatment, permeabilized cells were washed once with CSK buffer and treated with CSK buffer containing 0.1mg/mL RNase A (ThermoFisher) or RNase inhibitor for 5 min at 37°C. Washing with CSK buffer once again, cells were then fixed with 4% paraformaldehyde (PFA) in PBS for 10 min at room temperature. After fixation, cells were washed with PBS three times, stained with DAPI if required, and then imaged with microscope.

For DNase treatment, permeabilized cells were washed once with DNase digestion buffer (40 mM Tris-HCl pH 7.9, 10 mM NaCl, 6 mM MgCl2, 1mM CaCl2, 300 mM sucrose, 0.05% NP-40), and then incubated in DNase digestion buffer containing 100 units/mL DNase I (ThermoFisher) for 45 min at 37°C, twice for a total of 90 min. Lastly, nuclei were further extracted in CSK buffer supplemented with 0.25M ammonium sulfate for 15 min on ice to wash away digested chromatin, and fixed with 4% paraformaldehyde in PBS for 10 min at room temperature. After fixation, cells were washed with PBS three times, stained with DAPI if required, and then imaged with microscope.

### Auxin-induced degradation of SAF-A

For auxin-induced degradation of SAF-A in monoclonal homozygous SAF-A-mAID-Halo CMV-OsTIR1 HCT116 cells, indole-3-acetic acid (IAA, chemical analog of auxin) was treated in the cell growth media for 24 hr at 500μM from 1000× stock diluted in DMSO. Degradation was confirmed by the loss of fluorescence signal upon treatment of JF646 HaloTag to the cells.

### Acute depletion of SAF-A and mutant rescue

SNAP tagged mutant SAF-A containing Tet-On expression vectors were transfected with FuGene HD into monoclonal homozygous SAF-A-mAID-Halo CMV-OsTIR HCT116 cells. After 12 hr from transfection, 500μM of IAA and 100μM of doxycyclin were treated to the cells after replacement of cell growth media. At 36hr post-transfection, cells were treated with 250nM of JF646-HaloTag and 250nM of SNAP-Cell® Oregon Green (NEB Cat#S9104S) for 15 min at 37°C followed by washing with fresh cell growth media for 30 – 60 min at 37°C, and then imaged with microscope.

### EU labeling

Nascent RNAs in the nucleus were labeled using the Click-IT RNA Alexa Fluor Imaging Kit (ThermoFisher) with a modified protocol. Before pulse of 5-ethynyl uridine (EU), endogenous SAF-A-mAID-Halo HCT116 cells were labeled with 250nM of JF646-HaloTag for 15 min followed by washing with 37°C incubation of fresh cell growth media. We then exchanged the media with cell growth media containing 1mM of EU. Cells were incubated in the EU-containing media for 1 hr and fixed with 4% PFA. Click-labelling of EU incorporated into nascent RNA was performed according to the manufacturer’s protocol using AlexaFluor488 azide.

### Transcription inhibitor treatment

SAF-A-Dendra2 cells were plated on 12 well glass-bottom dishes (Cellvis, P12-1.5H-N) and maintained in growth media until reaching confluency of >50%. Before the treatment of transcription inhibitors, cells were incubated in the growth media with 250nM of JF646-HaloTag for 15 min to flourescently label the Pol II and washed briefly with PBS. JF646 treated cells were incubated at growth media with 0.1% DMSO (Sigma-Aldrich, D8418), 2μM actinomycin D (Sigma-Aldrich, A9415), or 100μM DRB (Sigma-Aldrich, D1916) diluted in DMSO for 6 hr. For DRB washout, DRB-treated cells were incubated in growth media without DRB for additional 6 hr.

### 7SK ASO transfection

Scramble (5’-mC*mG*mT*mC*G*A*T*G*T*G*A*T*G*C*T*G*T*mG*mT*mG*mA-3’) or 7SK anti-sense oligonucleotide (5’-mC*mC*mT*mT*G*A*G*A*G*C*T*T*G*T*T*T* G*mG*mA*mG*mG-3’) was transfected into cells with RNAiMAX (ThermoFisher Cat#13778075) or electroporation. In the case of the transfection with RNAiMAX, 30pmol of oligonucleotide was incubated in 300μl of Opti-MEM with 9μl of RNAiMAX for 5 min at room temperature. Then, 250μl of the mixture was treated to one million cells. All electroporations were performed using the 4D-Nucleofector™ X Unit and SE cell line kit (Lonza, Mt Waverley, VIC, Australia) with the concentration of 30pmol of oligonucleotide in the cuvette. After 24 hr from transfection, cells were treated with JF646-HaloTag followed by fixation and imaging.

### RT-qPCR

Total RNA was extracted from one million cells with a hybrid-R RNA prep kit (GeneAll Biotechnology Cat#305-101). RNA was reverse transcribed using M-MLV reverse transcriptase (Enzynomics). The resulting cDNA was used for RT-qPCR using the CFX-connect real-time PCR detection system (Bio-rad) for monitoring the synthesis of double-stranded DNA during PCR cycles using SYBR Green (Enzynomics). For each sample, triplicate test reactions were analyzed for the expression of the gene of interest, and the results were normalized to GAPDH mRNA.

### Nuclear speckle size analysis

For quantification of the nuclear speckle size on the 2D focal plane, images acquired from spinning-disk confocal microscopy (Nikon CSU-W1) were analyzed with Fiji software (23). In detail, the brightness of each cell image was normalized with a plugin ‘Local normalization’. Then, images were processed into the binary image to define the area occupied by the nuclear speckles. The area of the shapes in the binary image were measured as the sizes of the nuclear speckles, while shapes occupying area less than 0.1μm2 were regarded as noise and removed by the plugin ‘Shape filter’. SC35 were labeled with Alexa405 to acquire representative three-color images, whereas SC35 were labeled with Alexa647 to measure the size and the number of nuclear speckles.

### Super-resolution imaging

All super-resolution images in the study were acquired from a custom-built microscope based on the Nikon microscope body Ti2e with highly inclined and laminated optical (HILO) illumination. For PALM imaging of Dendra2, cells were illuminated with a 405-nm (for photoconversion of Dendra2) and a 561-nm laser (for excitation of photoconverted Dendra2). For each image, 4000 frames were acquired with the temporal resolution of 50 ms/frame. For dSTORM imaging of JF646-SAF-A-mAID-Halo, cells were excited with a 640-nm laser for 4000 frames with the temporal resolution of 50 ms/frame. For dSTORM imaging of Alexa Fluor488, optimal laser intensity was determined by excitation with a gradual increase of 488- nm laser intensity. Then the images were acquired with the temporal resolution of 20ms/frame for 45,000 frames in an imaging buffer containing 5% glucose, 100mM MEA, and 1% Glox solution (0.1% Glucose oxidase, 500U/ml Catalase in PBS) in PBS. In this study, lasers with a longer wavelength were illuminated prior to shorter ones, and flourophores with longer excitation wavelength were photobleached after imaging when needed, to minimize fluorescence cross-talk. Finally, collected images were analyzed and reconstructed into super-resolved images using a Fiji plugin ThunderSTORM and qSR^25^.

### Skeletonization analysis of the nuclear matrix

For quantitative measurements of the nuclear matrix, skeletonization was done with image processing software, Fiji, as described in fig. S10. In detail, super-resolved image of SAF-A was gaussian blurred and converted into binary image with local thresholding. Binary operations and shape filters were applied to smoothen the image and remove noises. Obtained binary images were converted into skeleton and quantitatively measured with ‘Skeletonize’ and ‘Analyze Skeleton’ plugins in Fiji^22^.

### Transcriptional condensate – nuclear matrix distance analysis

We identified transcriptional condensates on super-resolved images using the density-based spatial clustering of applications with noise (DBSCAN) algorithm^24^. Single molecule localizations of Pol II were grouped by DBSCAN, and each group defined as a single cluster, which corresponds to a transcriptional condensate. DBSCAN requires two parameters, the minimum number of detections in a group (N) and the minimum distance between detections in a group (R). In the study, we used N = 20 (points) and R = 30 (nm) to determine clusters.

Distance between the transcriptional condensates and the nuclear matrix was measured using center points of each transcriptional condensate and binary skeletonized image of the nuclear matrix. K-nearest neighbor algorithm with k=1 was applied to calculate the euclidean distance from each center point of transcriptional condensate to the nearest point of nuclear matrix.

### Immunofluorescence labeling

Cells were grown on 12 well glass-bottom dishes and fixed in 4% PFA as described above and stored at 4°C. Cells were permeabilized with 0.5% Triton X-100 (Sigma-Aldrich, T8787) in PBS for 10 min at RT. Following three washes in 0.1% Triton X-100 in PBS for 10 min, cells were blocked with blocking solution (1% Bovine Serum Albumin, 0.1% Triton X-100 in PBS) for 1 hr at RT and incubated with primary antibodies (anti-SC35 Abcam ab11826 1:200 dilution, anti-H3K27ac Abcam ab4729 1:200 dilution, anti-H3K27me3 Abcam ab6002 1:2000 dilution) diluted in blocking solution more than 16 hr at 4°C. After three washes in 0.1% Triton X-100 in PBS, the primary antibody was detected by secondary antibodies (Donkey Anti-Rabbit IgG H&L Alexa Fluor 647 Abcam ab150075 1:1000 dilution, Goat anti-Mouse IgG H&L Alexa Fluor 647 Abcam ab150115 1:1000 dilution, Donkey anti-Rabbit IgG H&L Alexa Fluor 405 Abcam ab175651 1:1000 dilution, Goat anti-Mouse IgG H&L Alexa Fluor 405 Abcam ab175660 1:1000 dilution) diluted in blocking solution at RT for 1 hr in the dark. Cells were washed three times with 0.1% Triton X-100 in PBS for 10 min.

### Whole chromosome painting

In situ hybridization of whole chromosome painting probes was performed following the protocol provided by the supplier (Creative Bioarray), with minor modifications. In detail, cells were fixed with 4% PFA in PBS for 10 min at room temperature. After fixation, cells were washed with PBS three times, and then permeabilized with 0.5% Tween-20 in PBS. Permeabilized cells were washed with PBS, and then dehydrated with 70% ethanol and stored overnight at 4°C. Next day, cells were rehydrated in cold PBS, followed by depurination with 0.1N HCl in room temperature for 5 min. Depurinated cells were dehydrated with serial incubation in 70%, 80%, 100% ethanol for 3 min each on ice. After dehydration, cells were air dried for ∼4 min. In the meantime, WCP probes were denatured in 80°C for 10 min, and incubated in 37°C water bath for 10-60 min before hybridization. Probes were protected from light during handling. Dehydrated cells were denatured with denaturation buffer (70% formamide in 2x saline sodium citrate (SSC) buffer pH adjusted to 7.0) in 80°C for 5 min. Appropriate time and temperature may differ for different cell types. Denatured cells were immediately dehydrated again by serial incubation in 70%, 80%, 100% ethanol for 3 min each on ice. After dehydration, cells were air dried for ∼4 minutes. WCP probes were applied directly on the dried cells, and covered with 18mm round coverslip. Then Cells and probes were co-denatured in 80°C for 4 min. After co-denaturation, the sample was incubated in dark, humidified chamber for hybridization in 37°C for 12-16 hr. After hybridization, the coverslip was removed, and the cells were washed with 0.1% Tween-20 in 0.4x SSC in 74°C for 4 min. Then the cells were washed again with 4x SSC in room temperature for 10 minutes. After washing, the cells were imaged in 2x SSC. When performed together with immunofluorescence labeling, immunofluorscence labeling was done priorly, and the sample was post-fixed with 4% PFA in PBS for 10 min at room temperature, and then the chromosome painting was performed as described above.

## Acknowledgments

This work was supported by the Suh Kyungbae Foundation (SUHF) and National Research Foundation (NRF) of Korea grants (2020R1C1C1014599 and 2020R1A4A3079755) to W.-K. Cho.

## Author contributions

G.P., K.R., and W.-K.C. conceived the project. G.P., K.R., and W.-K.C. organized the studies. H.S., K.R., C.K., J.L., Y.L., T.L.P., and G.P. contributed microscopy setup and cell culture. G.P., G.K., and J.L. generated cell lines. G.P., K.R., and G.K. developed and performed computational analyses. G.P. and K.R. analyzed and visualized all data in the study. G.P., K.R., and W.-K.C. wrote the manuscript. W.-K.C. supervised the project. G.P., K.R., T.L.P., and W.-K.C. contributed to editing the manuscript.

## Extended Data

**Extended Data Fig. S1.**
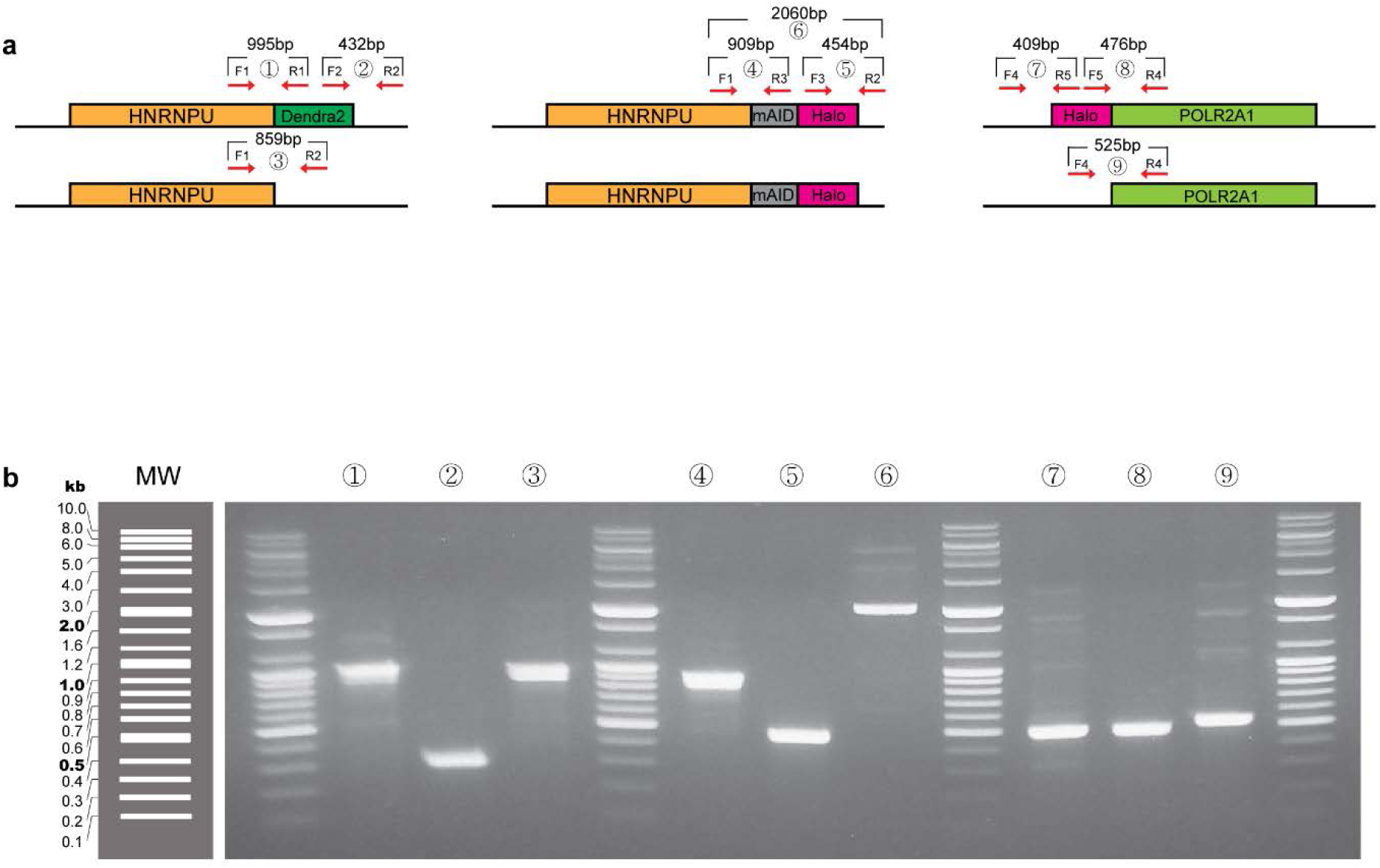
PCR validation for engineered cell lines by CRISPR/Cas9- knockin. (**a**) Schematic representation of genotyping for CRISPR/Cas9 engineered cell lines used for the study. Primers are listed in **Extended Data Table S1**. (**b**) Genotyping results for the cell lines after PCR amplification for target regions. Lane numbers denote primer pairs represented in (**a**).

**Extended Data Fig. S2.**
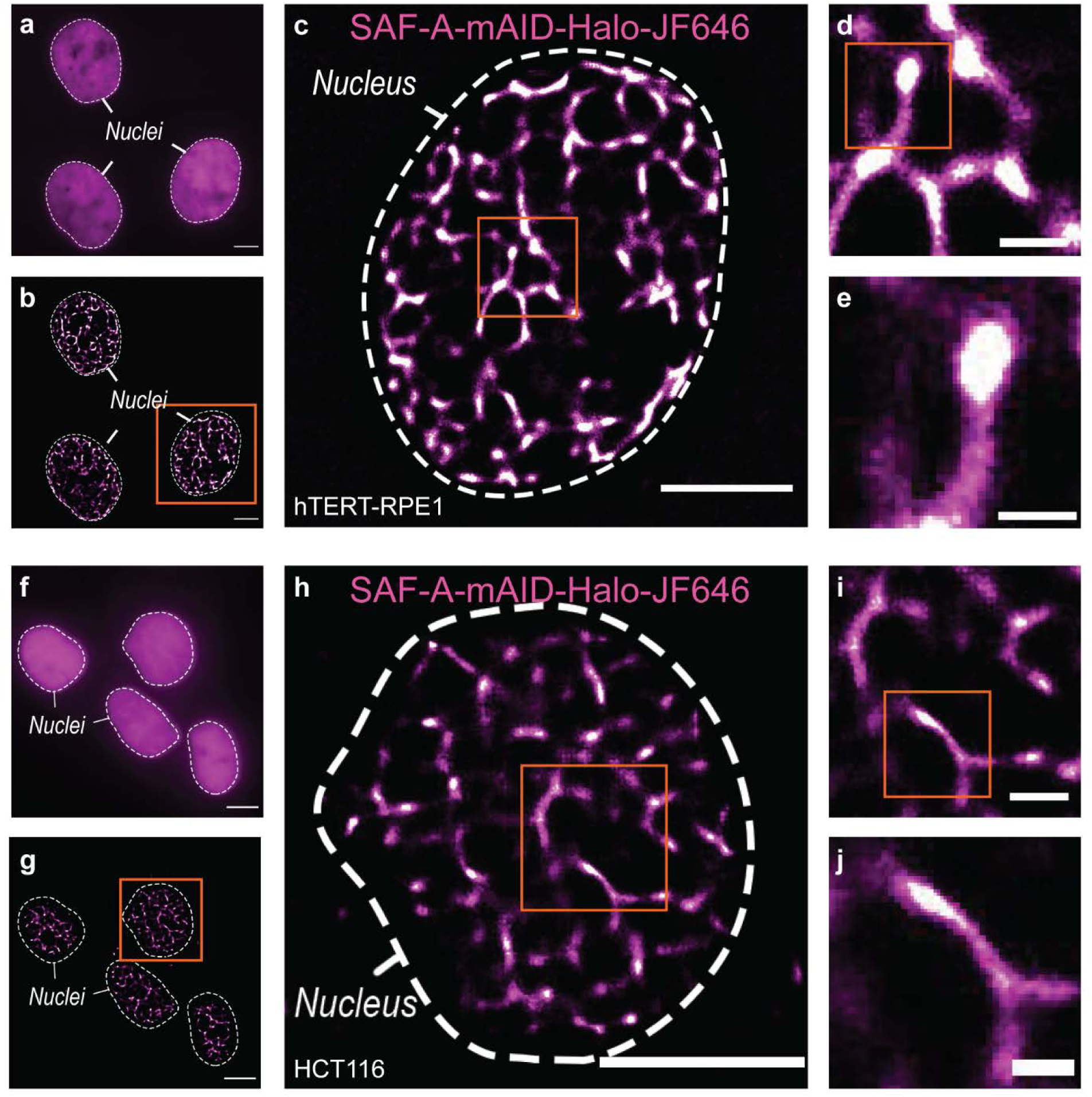
Super-resolution images of HaloTag-labeled SAF-A in hTERT-RPE1 and HCT116 cells. (**a and f**) A conventional fluorescence image of JF646 labeled SAF-A in fixed hTERT-RPE1 cells (**a**) and HCT116 cells (**f**). (**b and g**) Super-resolution (dSTORM) image of SAF-A. (**c to e**) Representative super-resolved images of SAF-A in a hTERT-RPE1 cell. Successive images are magnified versions of the boxed area in the previous image. (**h to j**) Representative super-resolved images of SAF-A in a HCT116 cell. Successive images are magnified versions of the boxed area in the previous image. JF646 labeled SAF-A via HaloTag also show the nuclear matrix structures. Scale bars represent 5 µm for entire nuclei images and 1 µm or 500 nm for zoom-in images.

**Extended Data Fig. S3.**
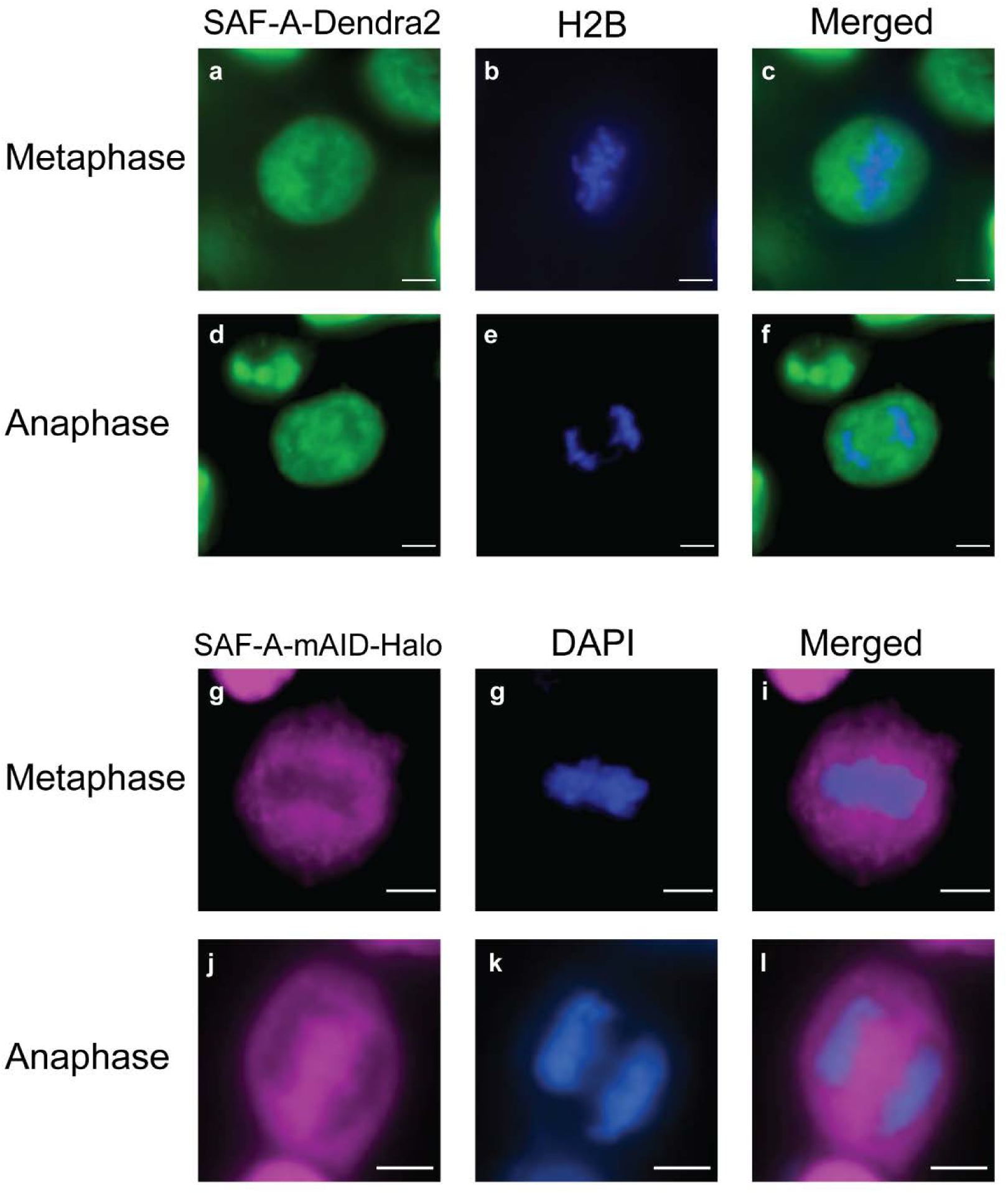
Exclusion of SAF-A from mitotic chromosomes. (**a to c**) Conventional fluorescence images for SAF-A-Dendra2 (**a**), H2B-EBFP2 (**b**) and a merged image (**c**) in metaphase. (**d to f**) Conventional fluorescence images for SAF-A-Dendra2 (**d**), H2B-EBFP2 (**e**) and a merged image (**f**) in anaphase. (**g to i**) Conventional fluorescence images for SAF-A-mAID-Halo-JF646 (**g**), DAPI staining (**h**) and a merged image **(i)** in metaphase. (**j to l**) Conventional fluorescence images for SAF-A-mAID-Halo-JF646 (**g**), DAPI staining (**h**) and a merged image (**i**) in anaphase. Scale bars, 5μm.

**Extended Data Fig. S4.**
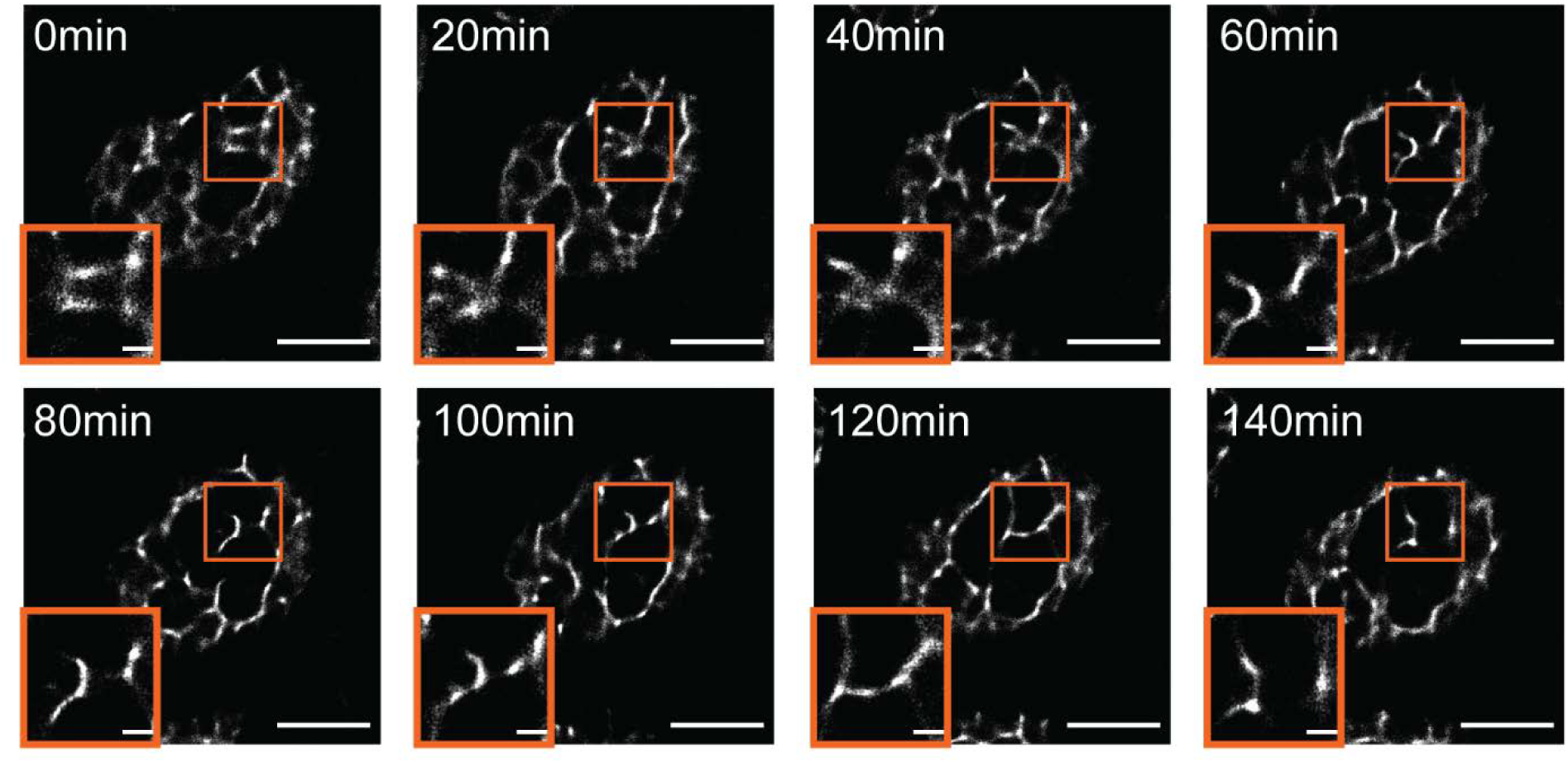
Dynamic structure of the nuclear matrix in live nuclei. Time-lapse super-resolved images of SAF-A-Dendra2 cells under DMSO treatment. Video available as **Extended Data Movie S2**. Scale bars, 5μm in overview, 1μm in zoom-in images.

**Extended Data Fig. S5.**
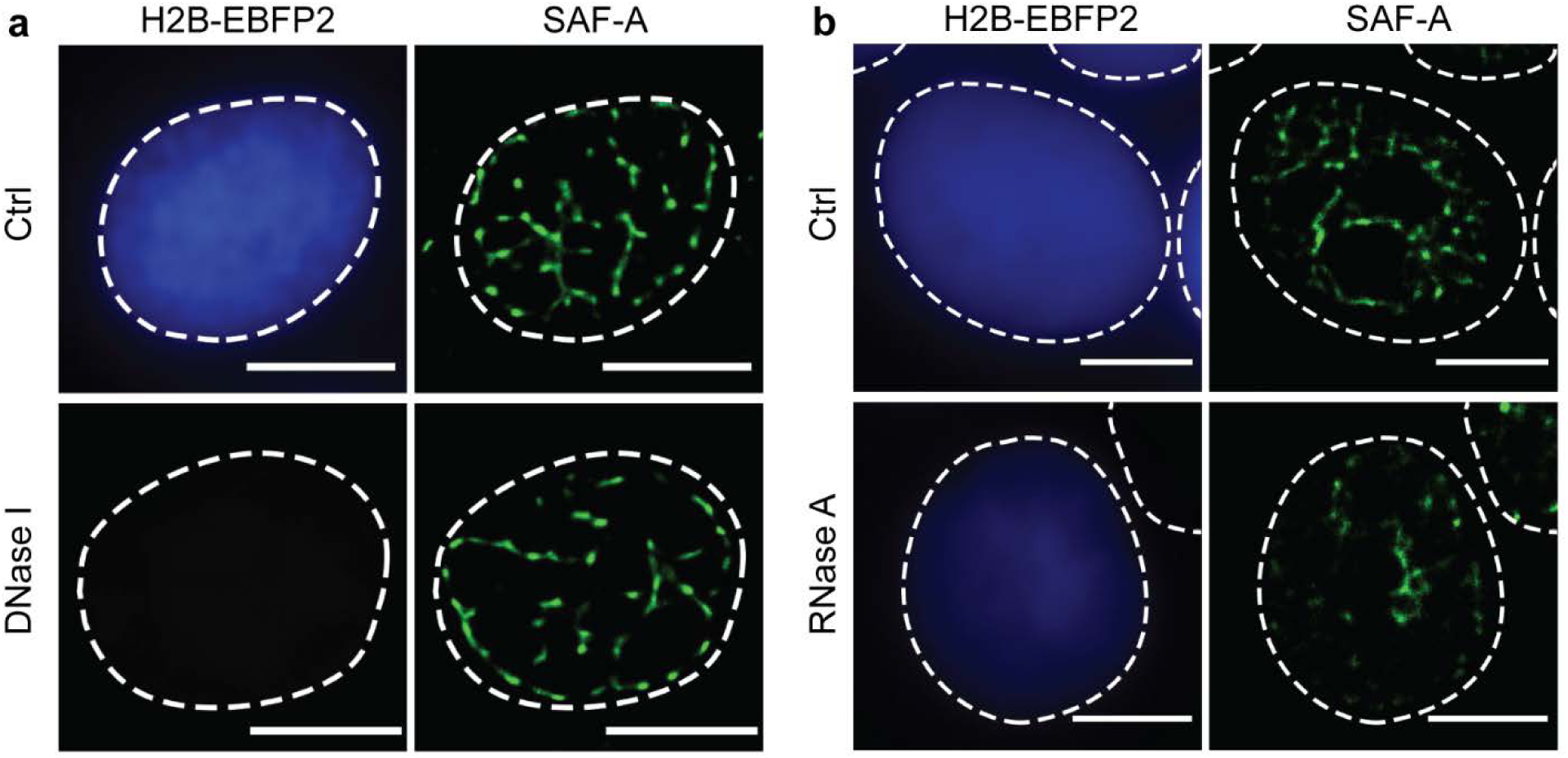
The nuclear matrix is RNA-dependent. Result of *in situ* fractionation followed by RNase or DNase treatment in HCT116 cells expressing H2B-EBFP2 and SAF-A-Dendra2. Results of (**a**) DNase I and (**b**) RNase A treatments. Scale bars, 5μm.

**Extended Data Fig. S6.**
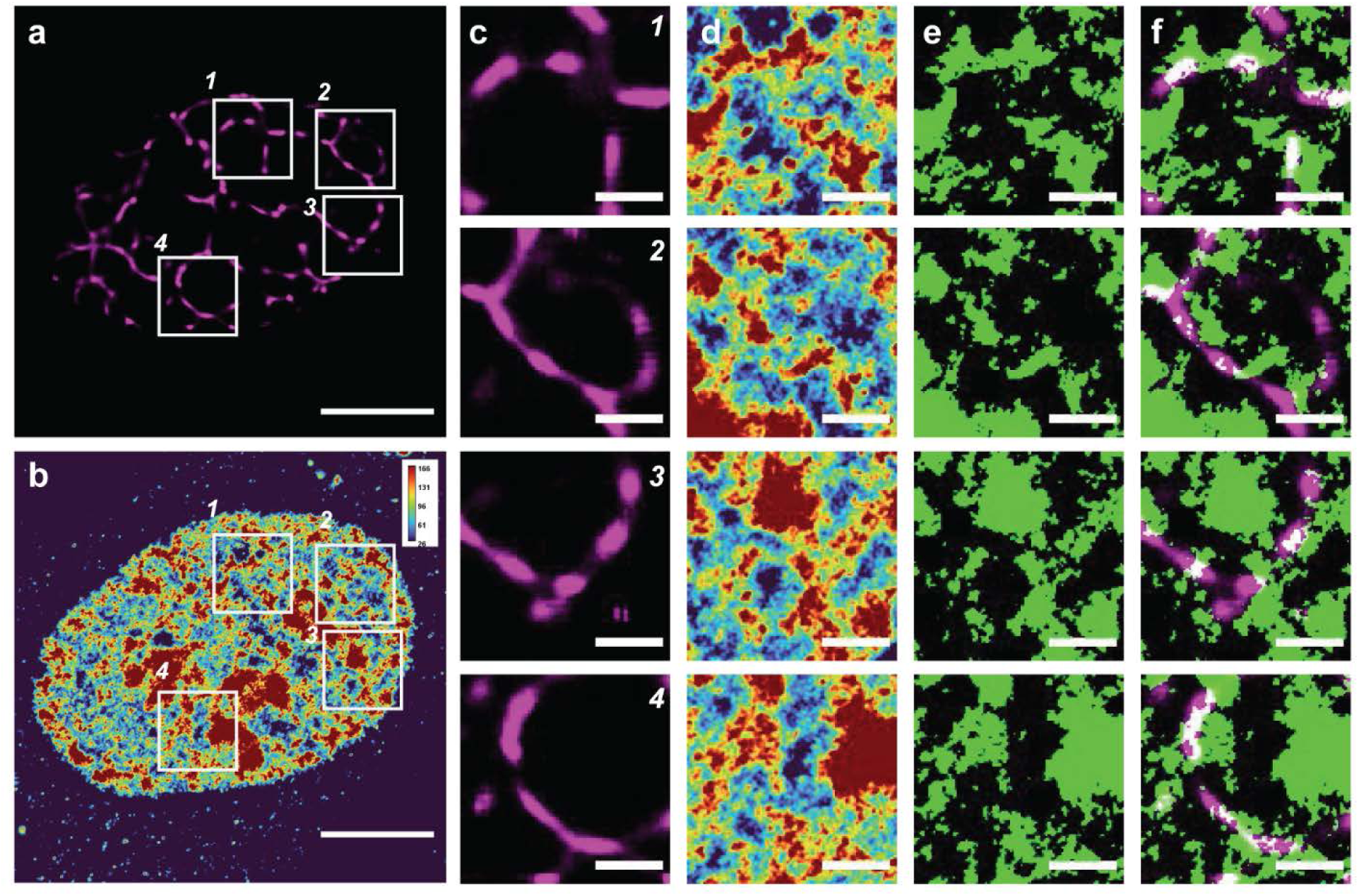
Capturing the nascent RNA enrichment near the nuclear matrix. The intranuclear localization of the nuclear matrix and the nascent RNA. (**a**) A super-resolved image (dSTORM) of JF646-HaloTag labeled SAF-A in a fixed HCT116 cell nucleus. (**b**) A corresponding image of density rendering for nuclear nascent RNA by EU labelling with Alexa Fluor 488. (**c**) Each numbered panel correspond to boxed area in (**a**). (**d**) Each numbered panels correspond to boxed area in (**b**). (**e**) Saturated images of intensive areas in (**d**). (**f**) Merged images of the super-resolved images of SAF-A (**c**) and nascent RNA enriched regions (**e**). Scale bars represent 5 µm for entire nuclei images and 1 µm for zoom-in images.

**Extended Data Fig. S7.**
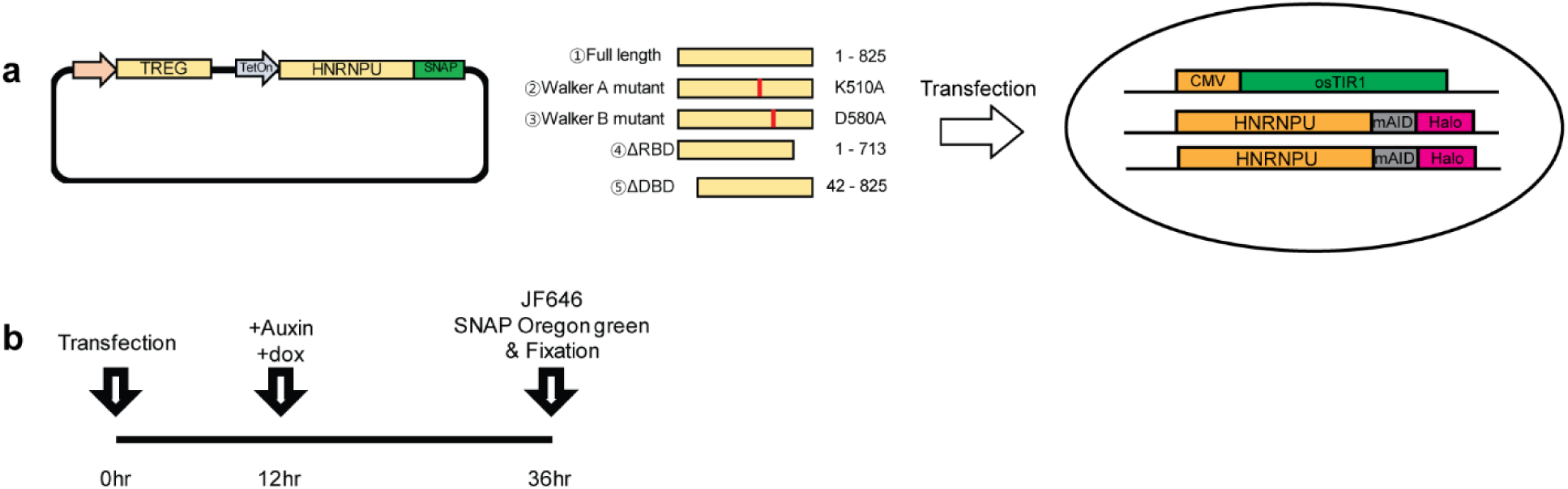
Schematic representation of an experimental procedure for acute depletion of SAF-A followed by overexpressing SAF-A mutant constructs. (**a**) Each SNAP-tagged SAF-A mutants were cloned in Tet-On expression vector and transiently transfected to the monoclonal homozygous SAF-A-mAID-Halo CMV-OsTIR cells. (**b**) The experimental time-line of the acute depletion of endogenous SAF-A followed by mutant rescue.

**Extended Data Fig. S8.**
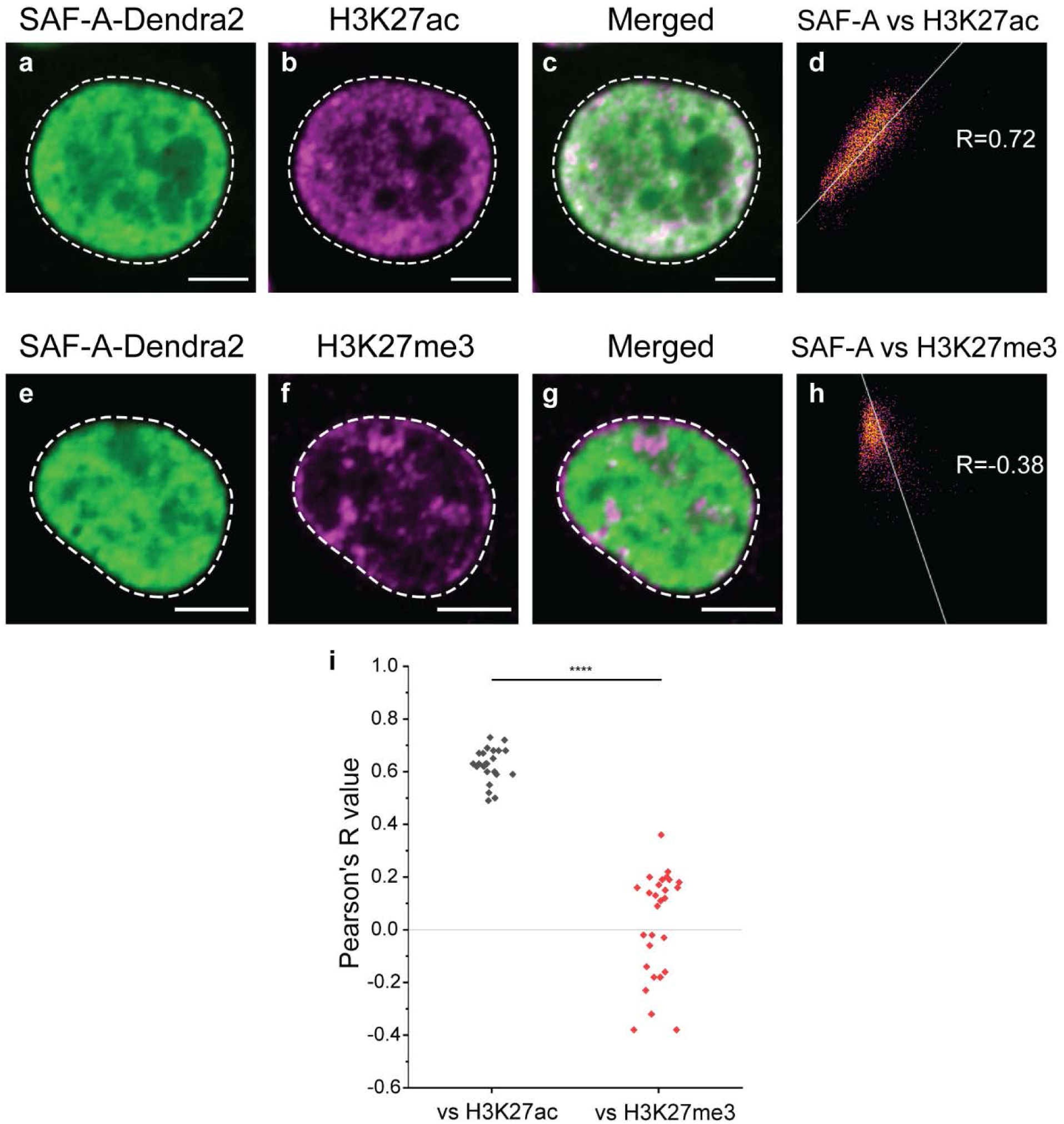
Dual-color fluorescence images of histone markers and SAF-A. (**a to c**) Dual-color images of SAF-A and a euchromatin histone marker, H3K27ac in a cell nucleus. (**d**) A dot plot for colocalization analysis between SAF-A and H3K27ac signals. (**e to g**) Dual-color images of SAF-A and a heterochromatin histone marker, H3K27me3 in a cell nucleus. (**h**) A dot plot for colocalization analysis between SAF-A and H3K27me3 signals. (**i**) Distributions of Pearson’s R values for SAF-A vs. H3K27ac and SAF-A vs. H3K27me3 from N = 22 cells and 27 cells, respectively. All images are obtained using spinning disk confocal microscopy (Nikon CSU-W1). Scale bars, 5μm.

**Extended Data Fig. S9.**
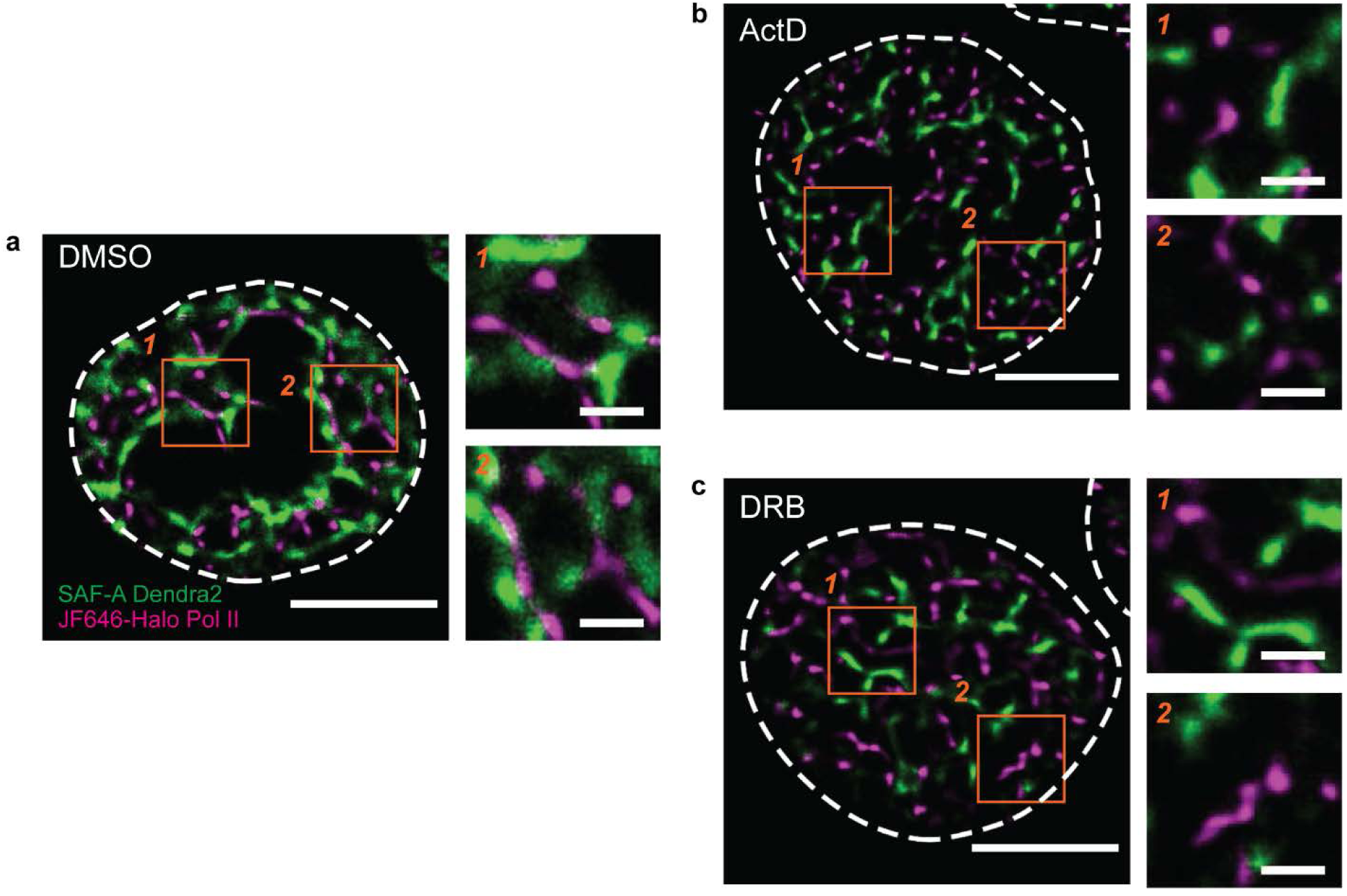
Dual-color imaging of SAF-A-Dendra2 and JF646-Halo Pol II in HCT116 cells after treatment of transcription inhibitors. Dual-color imaging of SAF-A-Dendra2 and JF646-Halo-Pol II after treatment of (**a**) DMSO, (**b**) ActD and (**c**) DRB. Each numbered panel corresponds to boxed areas in each image. Scale bars, 5μm in (**a**), (**b**) and (**c**), 1μm in zoom-in box.

**Extended Data Fig. S10.**
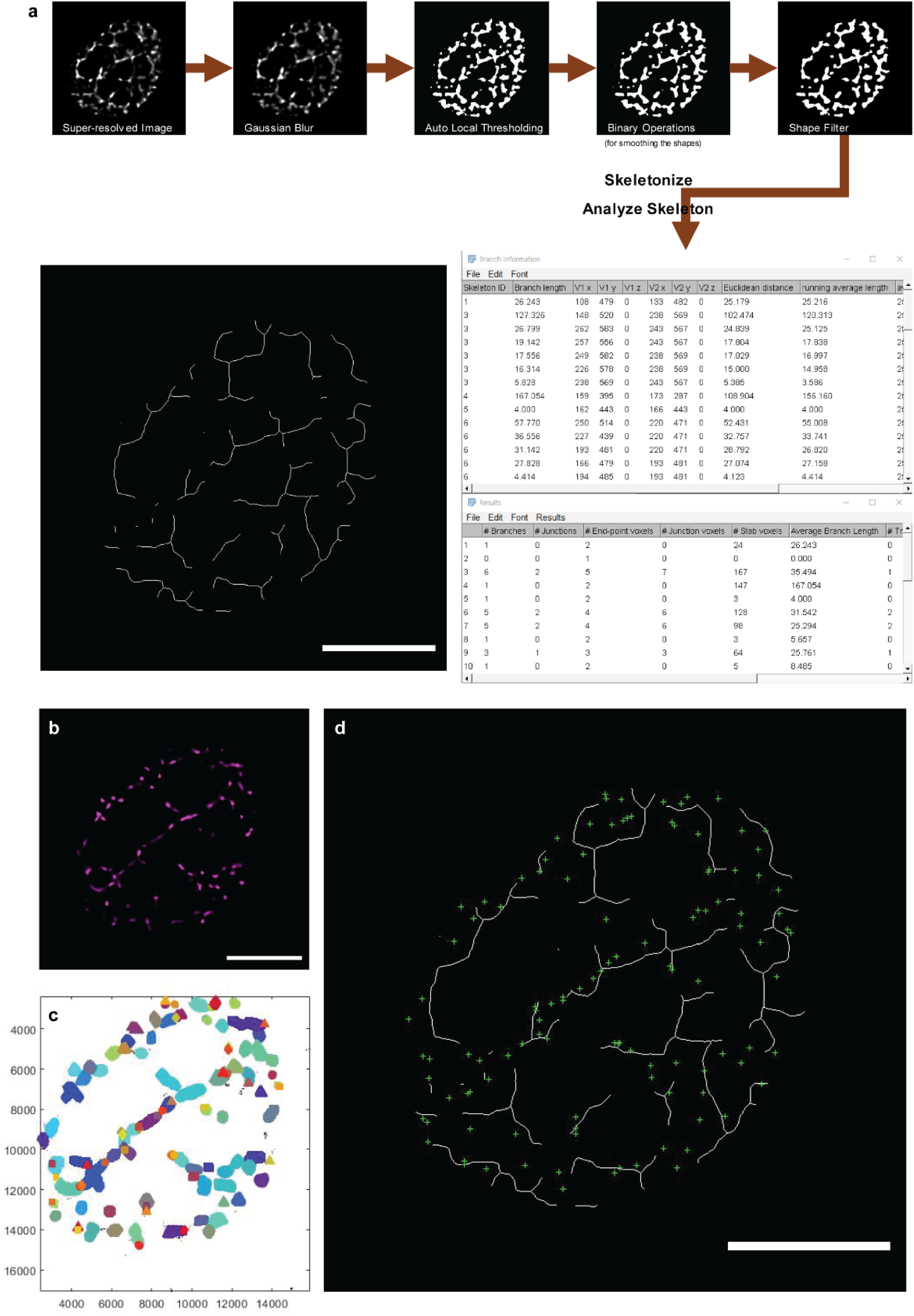
Schematic representation of quantitative measurement for the nuclear matrix and the transcriptional condensates. (**a**) Skeletonization process for the super-resolved images of nuclear matrix. (**b**) A super-resolved image of transcriptional condensates. (**c**) Defining transcriptional condensates by DBSCAN analysis for clustering. (**d**) A rendered image shows skeletonized nuclear matrix and center points of transcriptional condensates together. Scale bars, 5μm.

**Extended Data Fig. S11.**
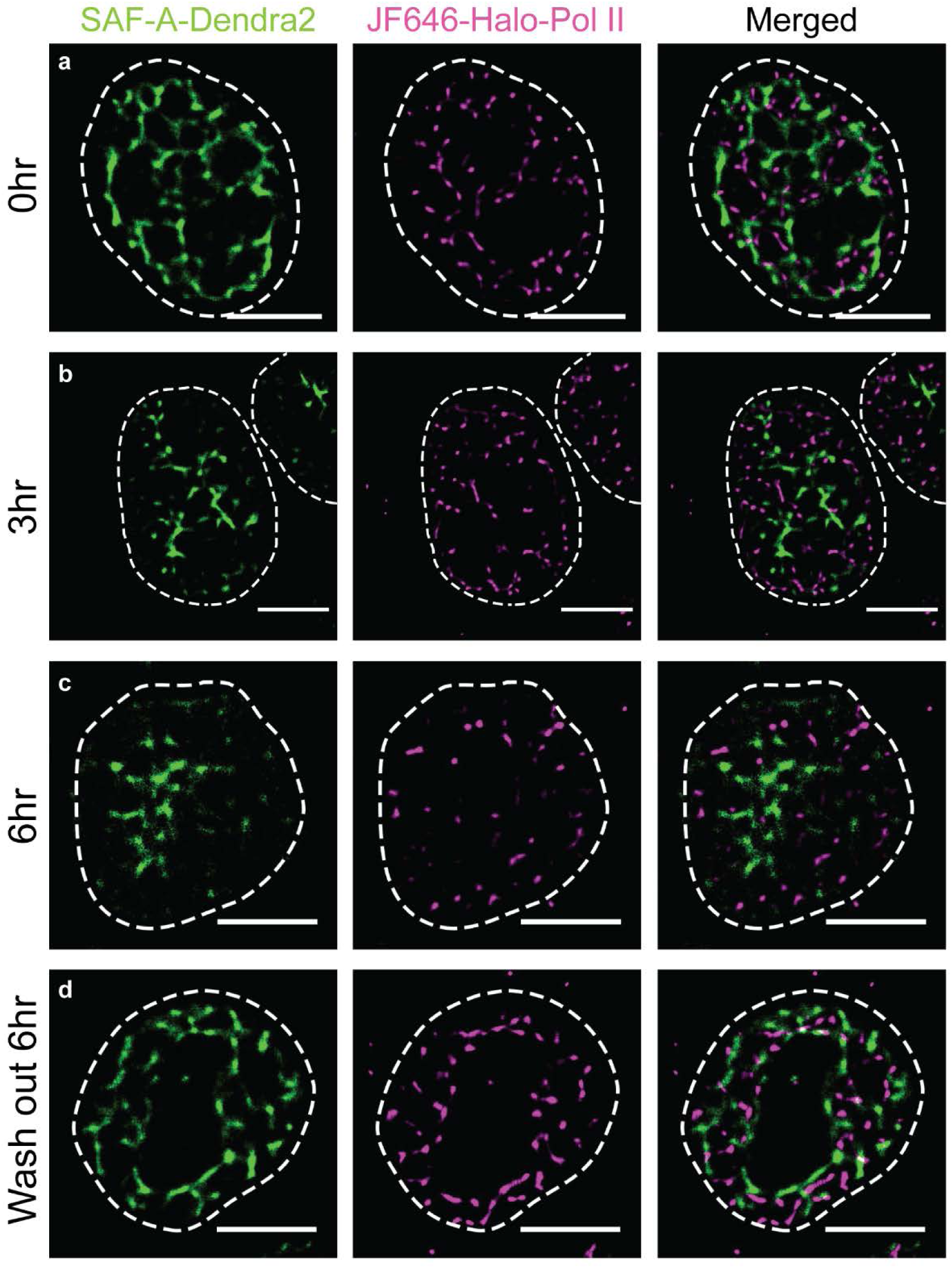
The disruption of the nuclear matrix upon transcription inhibition is dynamic and reversible. Disruption and restoration of the nuclear matrix and coordinated positioning of the transcriptional condensates upon DRB treatment and washout. Dual-color image of SAF-A- Dendra2 and JF646-Halo-Pol II in (**a**) untreated cell, (**b**) cell after 3 hours of DRB treatment, (**c**) cell after 6 hours of DRB treatment and (**d**) cell incubated without DRB for 6 hours, after 6 hours of DRB treatment. Scale bars, 5μm.

**Extended Data Fig. S12.**
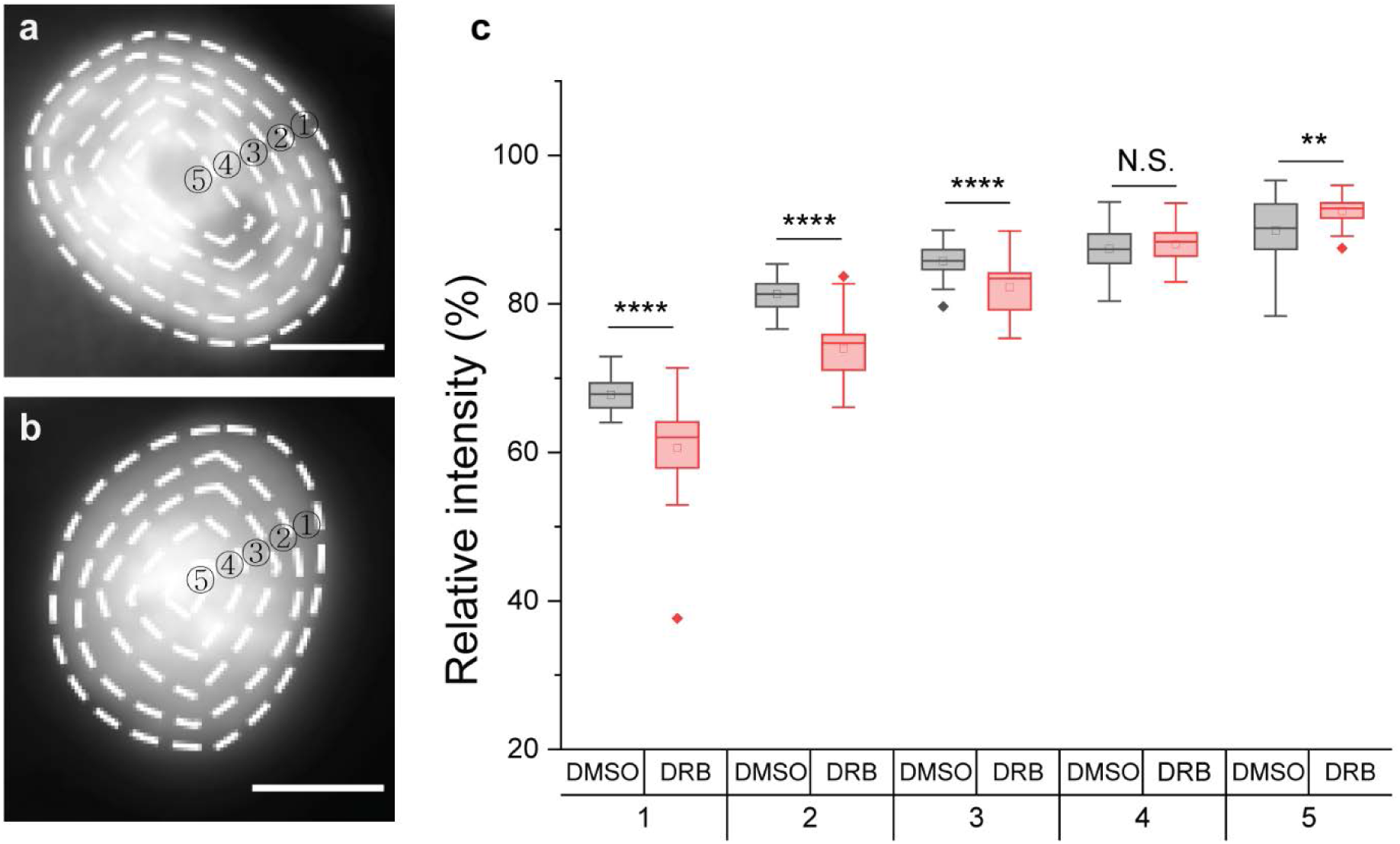
Radial distribution of SAF-A in a single nucleus changes upon transcription inhibition. The radial average intensity in a single nucleus of (**a**) DMSO or (**b**) DRB treated cell, respectively. Each nucleus is sectioned into five segments relative to the center of the nucleus. (**c**) Distribution of average intensity of each segment in drug-treated cells. (N = 30 cells from two experimental replicates) Error bars show mean ± standard deviation of the mean (s.d.m.). *, **, ***, **** indicate statistical significance at P < 0.05, P < 0.01, P < 0.001 and P < 0.0001 unpaired two-tailed Student’s t-tests. Scale bars, 5μm.

**Extended Data Fig. S13.**
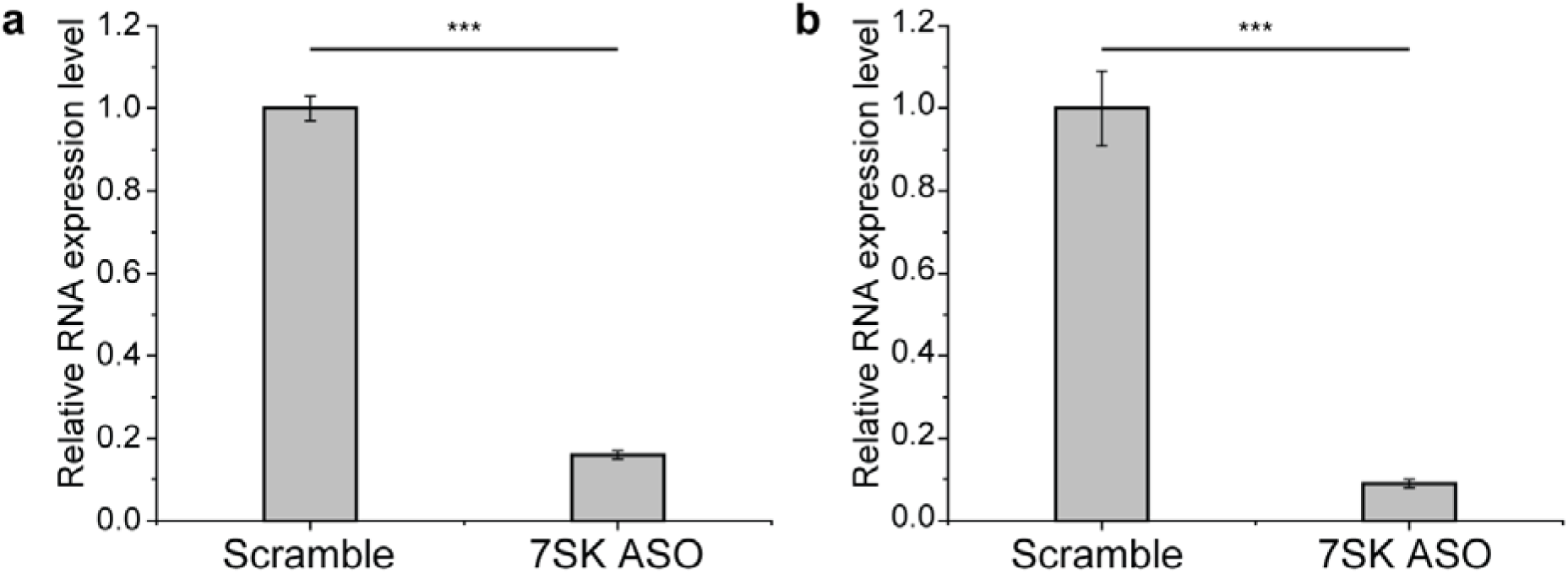
Validation of 7SK depletion by RT-qPCR in scramble or 7SK ASO transfected cells. Validation of 7SK depletion by RT-qPCR in scramble or 7SK ASO tranfected cells with (**a**) RNAiMAX or (**b**) nucleofection. (N = 3 for technical replicates) Error bars show mean ± standard deviation of the mean (s.d.m.). *, **, *** indicate statistical significance at P < 0.05, P < 0.01 and P < 0.001 unpaired two-tailed Student’s t-tests.

**Extended Data Fig. S14.**
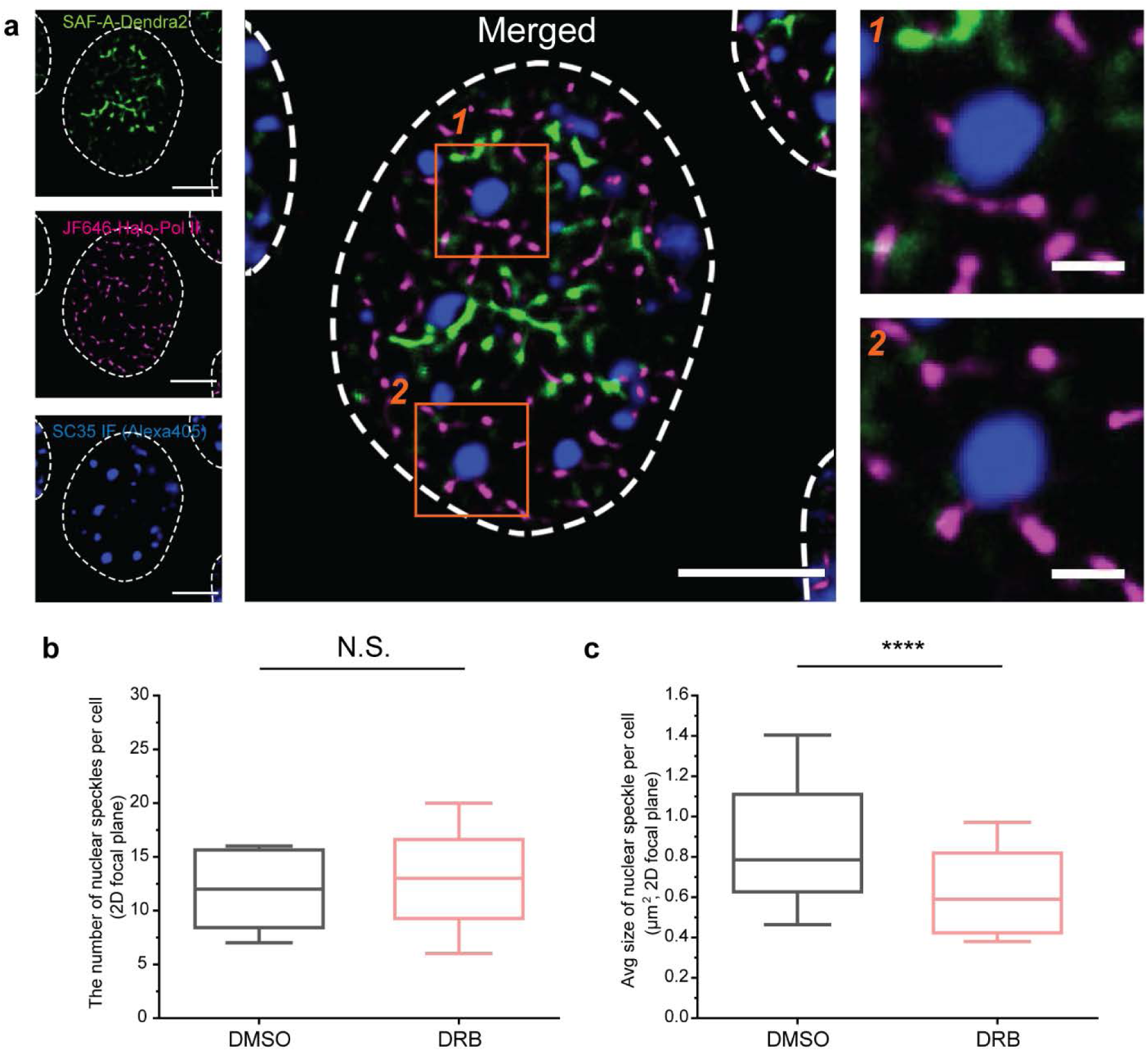
Disruption of the layered structure of transcriptional condensates and nuclear speckles relative to the nuclear matrix upon transcription inhibition by DRB treatment. (**a**) Three-color imaging of SAF-A-Dendra2, JF646-Halo-Pol II, and immunofluorescence of SC35 in DRB treated HCT116. (**b**) Distribution of the number of nuclear speckles in a single nucleus. (**c**) Distribution of the average size of nuclear speckles in a single nucleus. (N = 30 from two experimental replicates) Scale bars, 5μm in (**a**), 1μm in zoom-in box. Error bars show mean ± standard deviation of the mean (s.d.m.). *, **, ***, **** indicate statistical significance at P < 0.05, P < 0.01, P < 0.001 and P < 0.0001 unpaired two-tailed Student’s t-tests.

**Extended Data Fig. S15.**
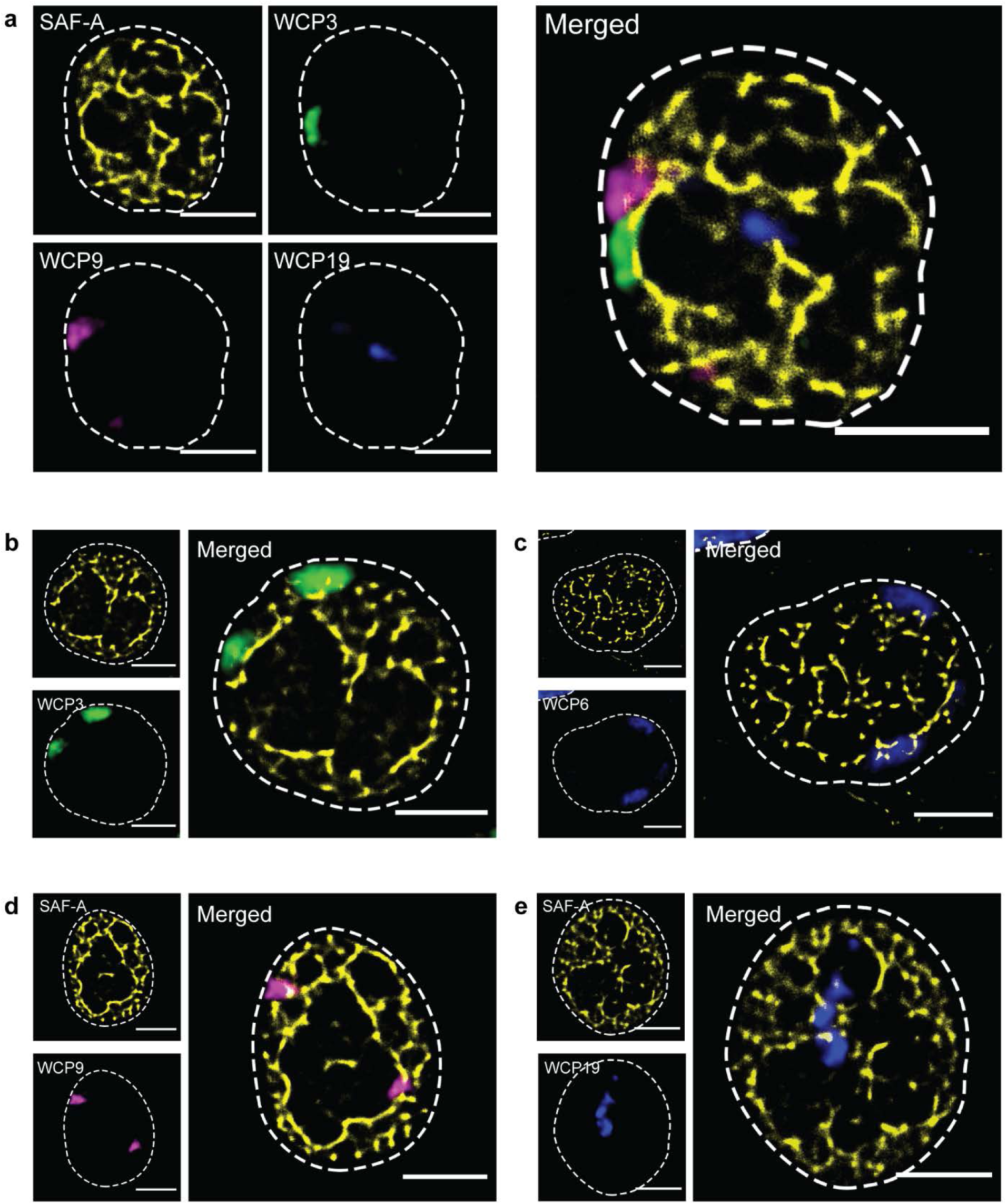
Visualization of chromosome territories relative to the nuclear matrix. (**a**) Multicolor image of chromosome 3, 9, and 19 visualized with whole chromosome painting, together with SAF-A-mAID-Halo-JF646. (**b-e**) Dual-color image of SAF-A-mAID-Halo-JF646 and (**b**) chromosome 3, (**c**) chromosome 6, (**d**) chromosome 9, (**e**) chromosome 19. Scale bars, 5μm.

**Extended Data Fig. S16.**
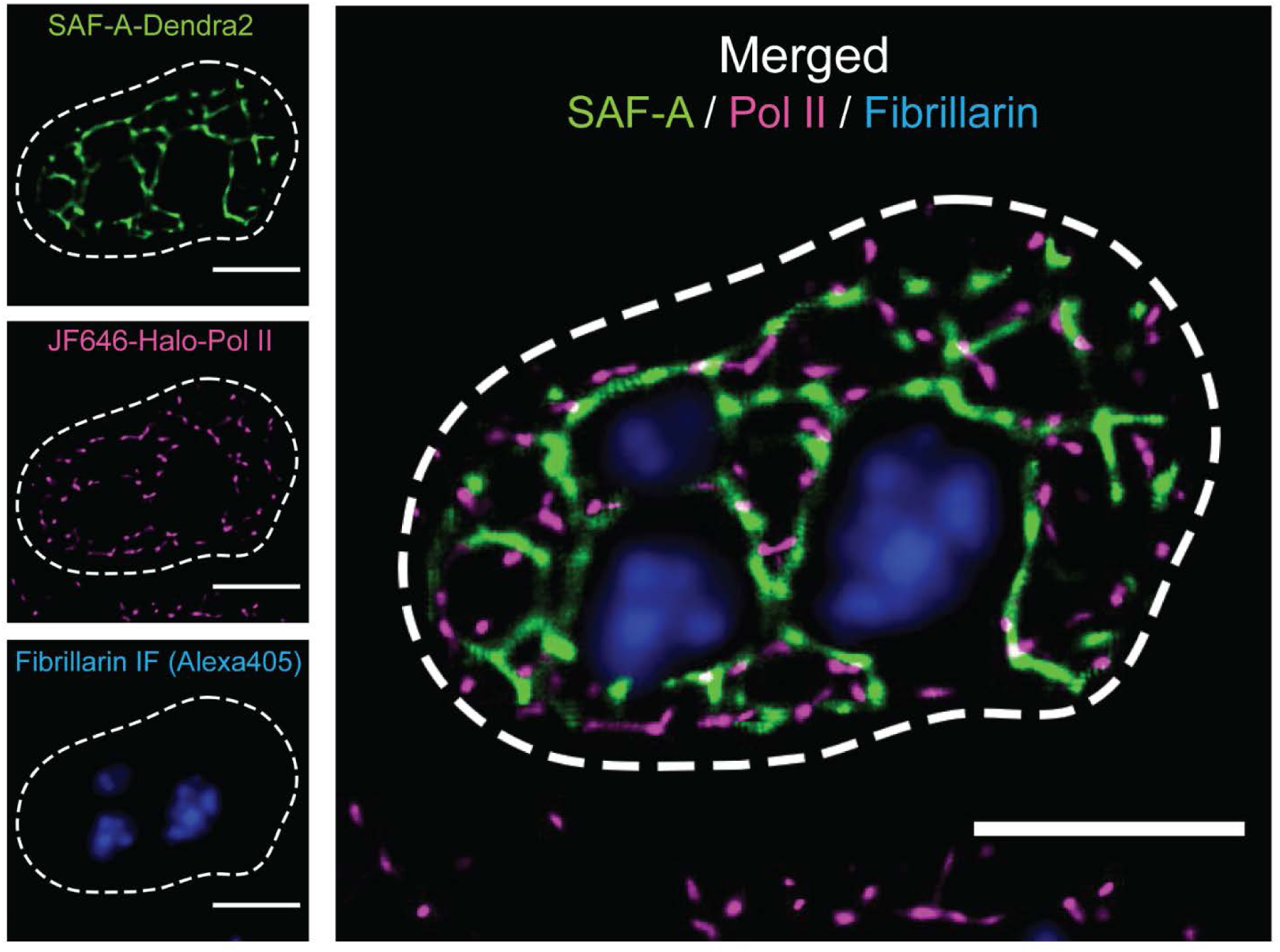
Visualization of the nucleoli along with transcriptional condensates and nuclear matrix by immunofluorescence labeling. Three-color image of SAF-A-Dendra2, JF646-Halo-Pol II and immunofluorescence of fibrillarin, nucleoli marker. Scale bar, 5μm.

**Extended Data Table S1.**
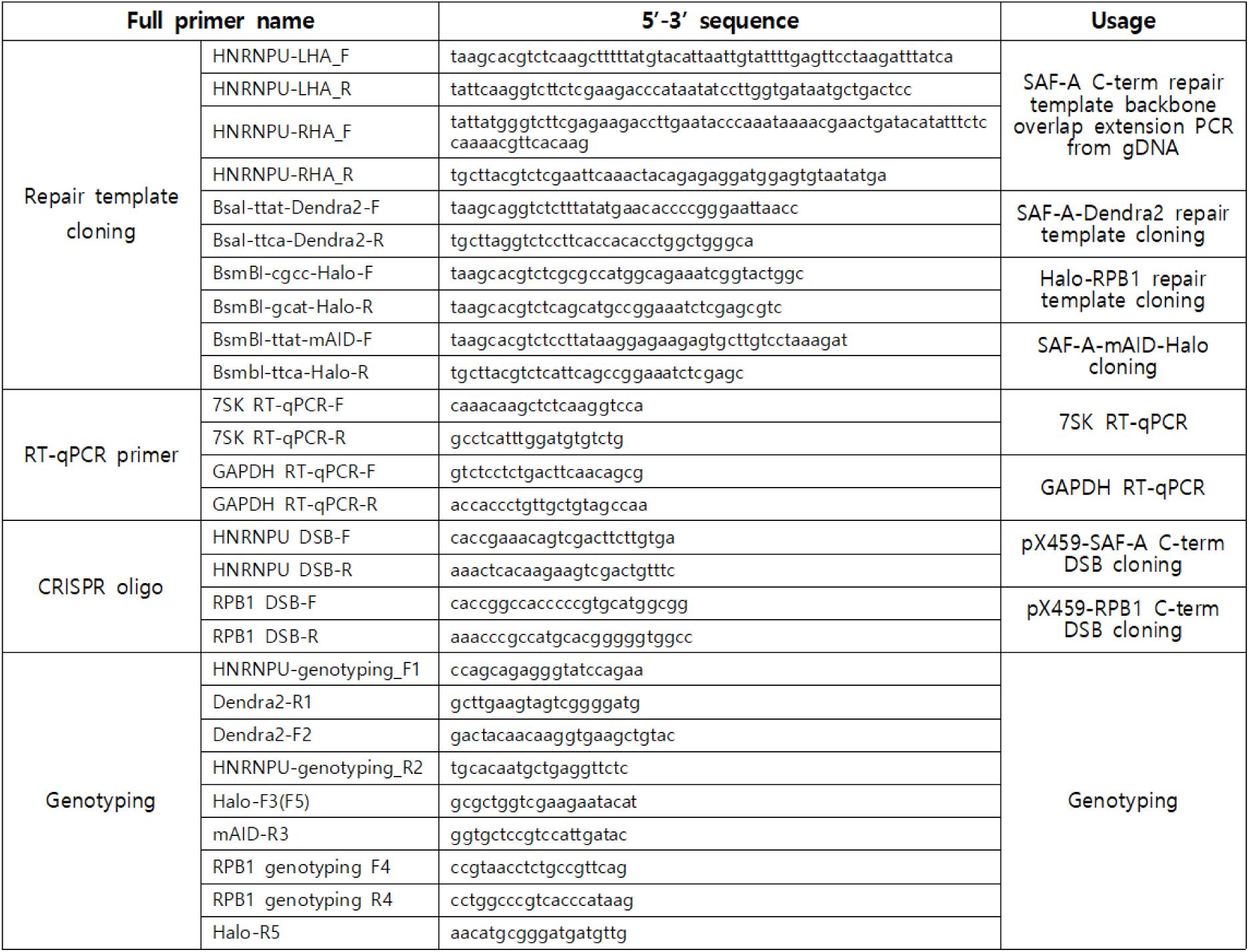
Oligo sequences for cloning and PCR.

**Extended Data Movie S1.**

The z-stack images of intranuclear SAF-A framework. Each image sequence was rendered from 500 frame images by ThunderSTORM with z-step size of 0.2μm. Scale bar, 5μm. The z-stack movements are repeated three times in the movie. The movie shows the structural network of the nuclear matrix extends throughout the entire nucleus.

**Extended Data Movie S2.**

The time-lapse movie of transformable structure of intranuclear SAF-A framework. Each image sequence was rendered from 500 frame images by ThunderSTORM with 20 min time interval. Scale bar, 10μm. The movie shows the nuclear matrix is dynamic in live cells.

